# The *Coxiella burnetii* QpH1 plasmid is a virulence factor for colonizing bone marrow-derived murine macrophages

**DOI:** 10.1101/2020.03.22.001875

**Authors:** Shengdong Luo, Zhihui Sun, Huahao Fan, Shanshan Lu, Yan Hu, Ruisheng Li, Xiaoping An, Yigang Tong, Lihua Song

## Abstract

*Coxiella burnetii* carries a large conserved plasmid or plasmid-like chromosomally integrated sequence of unknown function. Here we report the curing of QpH1 plasmid from *C. burnetii* Nine Mile phase II, the characterization of QpH1-deficient *C. burnetii* in in vitro and in vivo infection models, and the characterization of plasmid biology. A shuttle vector pQGK, which is composed of the CBUA0036-0039a region (predicted for QpH1 maintenance), an *E. coli* plasmid ori, eGFP and kanamycin resistance genes was constructed. The pQGK vector can be stably transformed into Nine Mile II and maintained at a similar low copy like QpH1. Importantly, transformation with pQGK cured the endogenous QpH1 due to plasmid incompatibility. Compared to a Nine Mile II transformant of a RSF1010-based vector, the pQGK transformant shows an identical one-step growth curve in axenic media, a similar growth curve in Buffalo green monkey kidney cells, an evident growth defect in macrophage-like THP-1 cells, and dramatically reduced ability of colonizing bone marrow-derived murine macrophages. In the SCID mouse infection model, the pQGK transformants caused a lesser extent of splenomegaly. Moreover, the plasmid biology was investigated by mutagenesis. We found CBUA0037-0039 are essential for plasmid maintenance, and CBUA0037-0038 account for plasmid compatibility. Taken together, our data suggest that QpH1 encodes factor(s) essential for colonizing murine macrophages, and to a lesser extent for colonizing human macrophages. This study highlights a critical role of QpH1 for *C. burnetii* persistence in rodents, and expands the toolkit for genetic studies in *C. burnetii*.

**Author summary:** It is postulated that *C. burnetii* recently evolved from an inherited symbiont of ticks by the acquisition of novel virulence factors. All *C. burnetii* isolates carry a large plasmid or have a chromosomally integrated plasmid-like sequence. The plasmid is a candidate virulence factor that contributes to *C. burnetii* evolution. Here we describe the construction of novel shuttle vectors that allow to make plasmid-deficient *C. burnetii* mutants. With this plasmid-curing approach, we characterized the role of the QpH1 plasmid in *in vitro* and *in vivo C. burnetii* infection models. We found that the plasmid plays a critical role for *C. burnetii* growth in macrophages, especially in murine macrophages, but not in axenic media and BGMK cells. Our work highlights an essential role of the plasmid for the acquisition of colonizing capability in rodents by *C. burnetii*. This study represents a major step toward unravelling the mystery of the *C. burnetii* cryptic plasmids.

## Introduction

*C. burnetii* is a Gram-negative intracellular bacterium that causes Q fever, a world widely distributed zoonosis [1]. It is highly infectious and is classified as a potential biowarfare agent [2]. Its infections in humans are mostly asymptomatic but may manifest as an acute disease (mainly as a self-limiting febrile illness, pneumonia, or hepatitis) or as a chronic disease (mainly endocarditis in patients with previous valvulopathy) [1]. All *C. burnetii* isolates maintain a large cryptic plasmid or plasmid-like chromosomally integrated sequence [3]. This absolute conservation of plasmid sequences suggests they are critical in *C. burnetii* survival [4]. Increased knowledge of the plasmid sequences will enhance our understanding of *C. burnetii* biology and pathogenesis and will be important for guiding the design of live attenuated vaccine strains for the prevention of Q fever.

*C. burnetii* has at least five different plasmid types: four different plasmids (QpH1, QpRS, QpDV, and QpDG) and one type of QpRS-like chromosomally integrated sequence [5–9]. The plasmids range from 32 to 54 kb in size and share a 25 kb core region [3]. Characterization of these plasmid types led to the classification of five associated genomic groups. The hypothesized correlation of plasmid types and genomic groups with the clinical outcomes of Q fever was controversial [3, 10, 11], but the concept of genomic-group specific virulence was supported by mouse and guinea pig infection studies [12, 13]. According to studies on *C. burnetii* mostly from Europe and North America, this bacterium is considered having a clonal population structure with low genetic diversity [14–16]. Future studies on more *C. burnetii* isolates especially from other regions will help to understand its global genetic diversity.

The sequences of all five representative plasmid types in *C. burnetii* have been determined [3, 7, 8, 17, 18]. Paul et al. analyzed the diversity of plasmid genes by using DNA microarrays [3]. Nine predicted functional plasmid ORFs are conserved in all plasmid sequences. Two of these ORFs appear to encode proteins similar to phage proteins involved in tail assembly and site-specific recombination, while seven of them encode hypothetical proteins. Three (CBUA0006, -0013 and -0023) of these conserved plasmid hypothetical proteins are secreted effectors of the type IV secretion system [4]. During ectopic expression in HeLa cells, these effectors localize at different subcellular sites, suggesting roles in subversion of host cell functions. In addition, novel plasmid-specific effectors were also identified, supporting the concept of pathotype virulence [19]. Besides the identification of plasmid-encoded secretion effectors, the plasmid is also speculated to be able to be transferred to the host cell [20].

The advent and foregoing improvement of axenic culture techniques greatly facilitates studies on *C. burnetii* biology [21–24]. *C. burnetii* replicates within phagolysosome-like parasitophorous vacuoles (termed *Coxiella*-containing vacuole, or CCV) in mononuclear phagocytes. Genetic studies revealed that the CCV biogenesis and *C. burnetii* growth in host cells require a type IV secretion system and a repertoire of type IV effectors [25–28]. The role that these effectors play in *C. burnetii* pathogenicity is of intense interest. However, regarding the plasmid effectors, reports on their mutants are scarce. By using transposon mutagenesis, Martinez et al. identified mutants of eight QpH1 genes and observed strong phenotypes with mutants of *parB* and *repA*, surprisingly, not the mutant of the conserved secretion effector CBUA0023 [29]. Regardless of the identification of plasmid secretion effectors, a role for the plasmid in *C. burnetii* pathogenesis remains elusive.

Since the first development of axenic media, the *C. burnetii* genetics toolbox has been expanding [30, 31]. Current genetic tools for *C. burnetii* include, but are not limited to, transposon systems [32], RSF1010 ori-based shuttle vectors [4, 33] and targeted gene inactivation systems [31]. However, there have been no reports of studies using these tools to decipher roles of plasmids in *C. burnetii* pathogenesis. Here, we describe a set of QpH1-derived shuttle vectors for *C. burnetii* transformation. These shuttle vectors have different compatibilities with QpH1 and different stabilities in *C. burnetii*. A QpH1-deficient strain of *C. burnetii* was constructed by transforming with the pQGK vector. Infection experiments show that QpH1 is essential for colonizing bone marrow-derived murine macrophages. This work increases our knowledge on plasmid biology and provides new genetic tools for investigating *C. burnetii* pathogenesis.

## Results

### *C. burnetii* plasmid has a putative iteron-like origin of replication

The QpH1 plasmid of the Nine Mile isolate is 37,319-bp in size and encodes 40 ORFs, of which only 11 are similar to proteins of known function (Figure 1A) [18]. Besides its cryptic role in pathogenesis, the plasmid’s mechanism of replication and partitioning also lacks investigation. A gene cluster CBUA0036-39 encodes homologs of ParB, ParA, RepB and RepA, respectively, which are supposed to function in plasmid replication and partitioning based on protein similarities. The plasmid was found to have a low copy number [10], but its origin of replication has never been identified. With equicktandem of the EMBOSS software suite [34, 35], we found a putative origin of replication -a region composed of four 21-bp iteron-like sequences between CBUA0036 and -37 (Figure 1B). Unlike the highly conserved iterons in other plasmids, all four iteron-like sequences in QpH1 have base variations from each other. And in currently known *C. burnetii* plasmids, this putative iteron region is highly conserved. The finding of this putative iteron region suggests genes required for QpH1 replication and partitioning are clustered.

**Figure 1.**
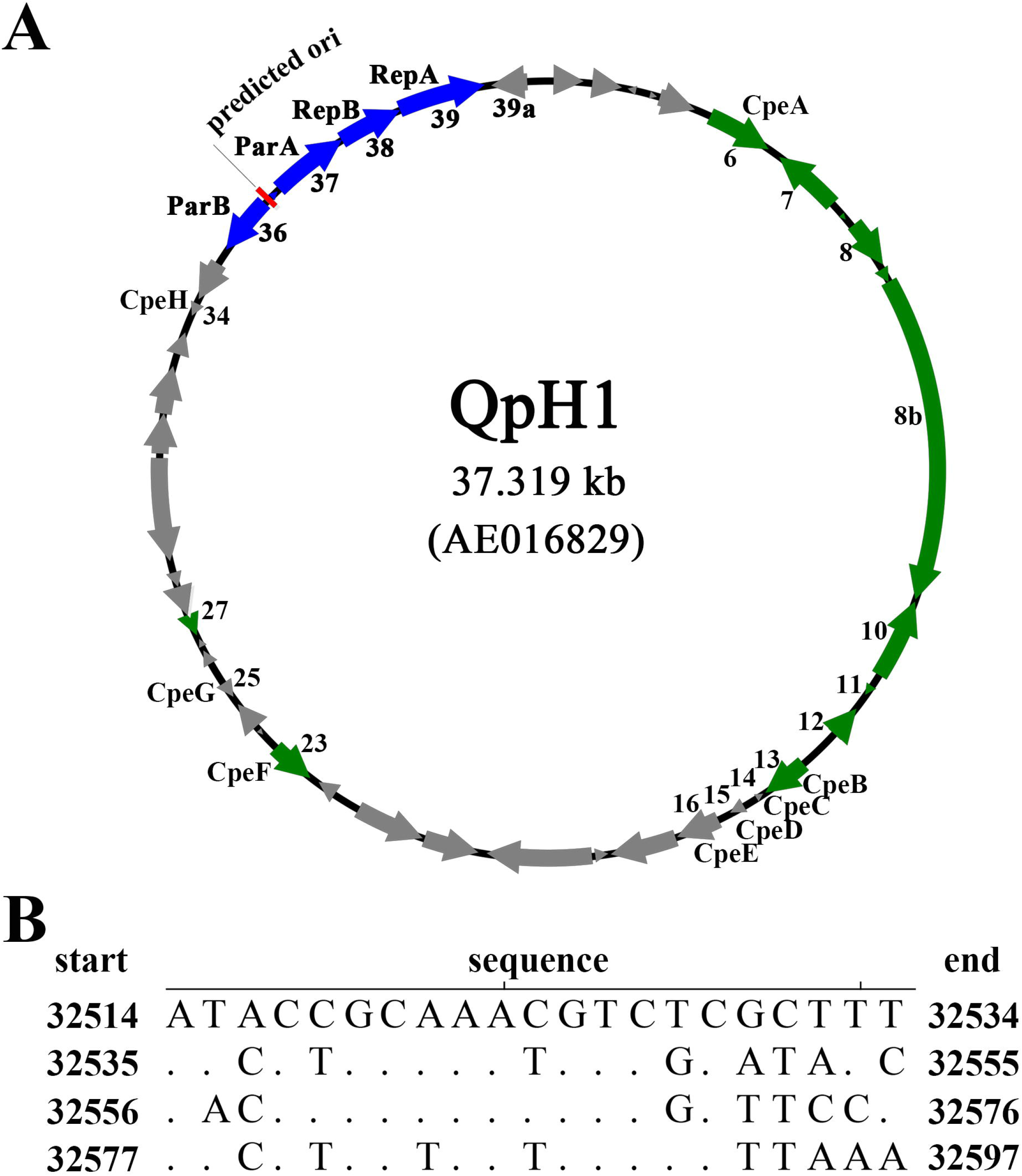
Map of the QpH1 plasmid (A) and the sequence of its predicted origin of replication, which is composed of four 21-bp iteron-like sequences (B). The conserved ORFs (green) [3], predicted ORFs for plasmid maintenance (blue) and predicted ori (red) are as indicated (A). In the iteron alignment (B), dots represent nucleotides that are identical to those in the most upper iteron. The first and last nucleotides in each iteron are given as its position in the complete plasmid.

### Construction of incompatible plasmid pQGK and curing of endogenous QpH1 from Nine Mile II by transformation

The RSF1010-ori based shuttle vectors are routinely used for *C. burnetii* transformation [31, 33]. We first made a RSF1010-ori based shuttle vector pMMGK for transformation (Figure 2A). An incompatible plasmid pQGK was then constructed as suggested by the clustering of genes (CBUA0036-39) for QpH1 maintenance (Figure 2B). The hypothetical CBUA0039a gene was also incorporated in pQGK, as this gene locates close to CBUA0039 and might play unknown roles in plasmid maintenance. The plasmids -pMMGK and pQGK were transformed into Nine Mile II, respectively. After passaging in the presence of kanamycin for six to seven times, the transformants were cloned by using semi-solid ACCM-2 plates and characterized by PCR detection of QpH1 ORFs. All QpH1 ORFs can be detected in NMII*pMMGK*, but only the CBUA0036-39a ORFs can be detected in NMII*pQGK* (Figure 3). The absence of CBUA001-0034a in NMII*pQGK* indicates that the endogenous QpH1 was cured by transformation of pQGK.

**Figure 2.**
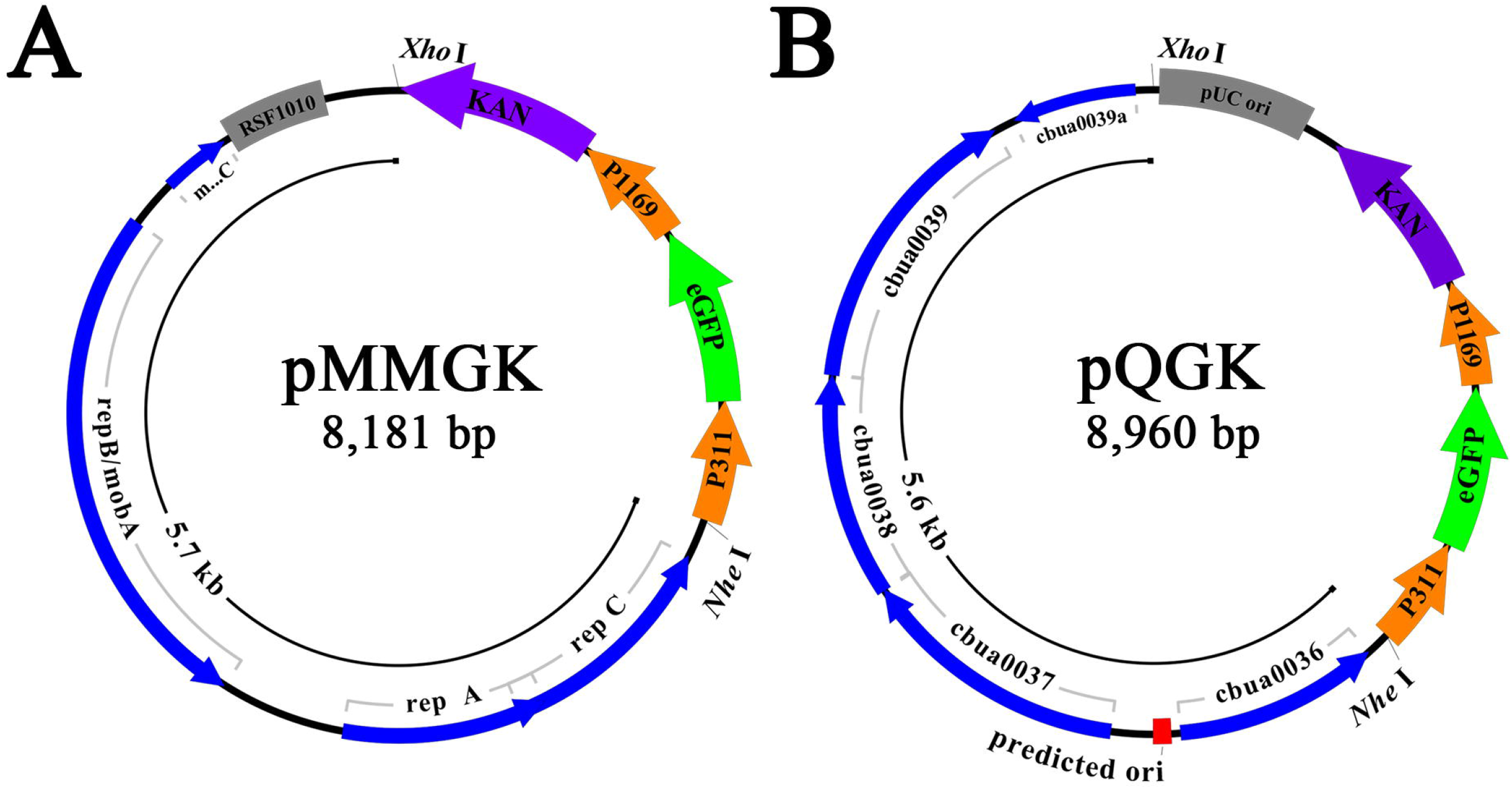
Map of two shuttle vectors -pMMGK (A) and pQGK (B). Both vectors share identical Kan^R^ and eGFP genes, driven by P1169 and P311 promotors, respectively. The inner circle of the pMMGK vector represents the backbone of a RSF1010 ori-based plasmid pMMB207 (A). The inner circle of the pQGK vector represents the fragment from QpH1 (B). The genes and their direction of transcription are represented by the boxes of the outer circle.

**Figure 3.**
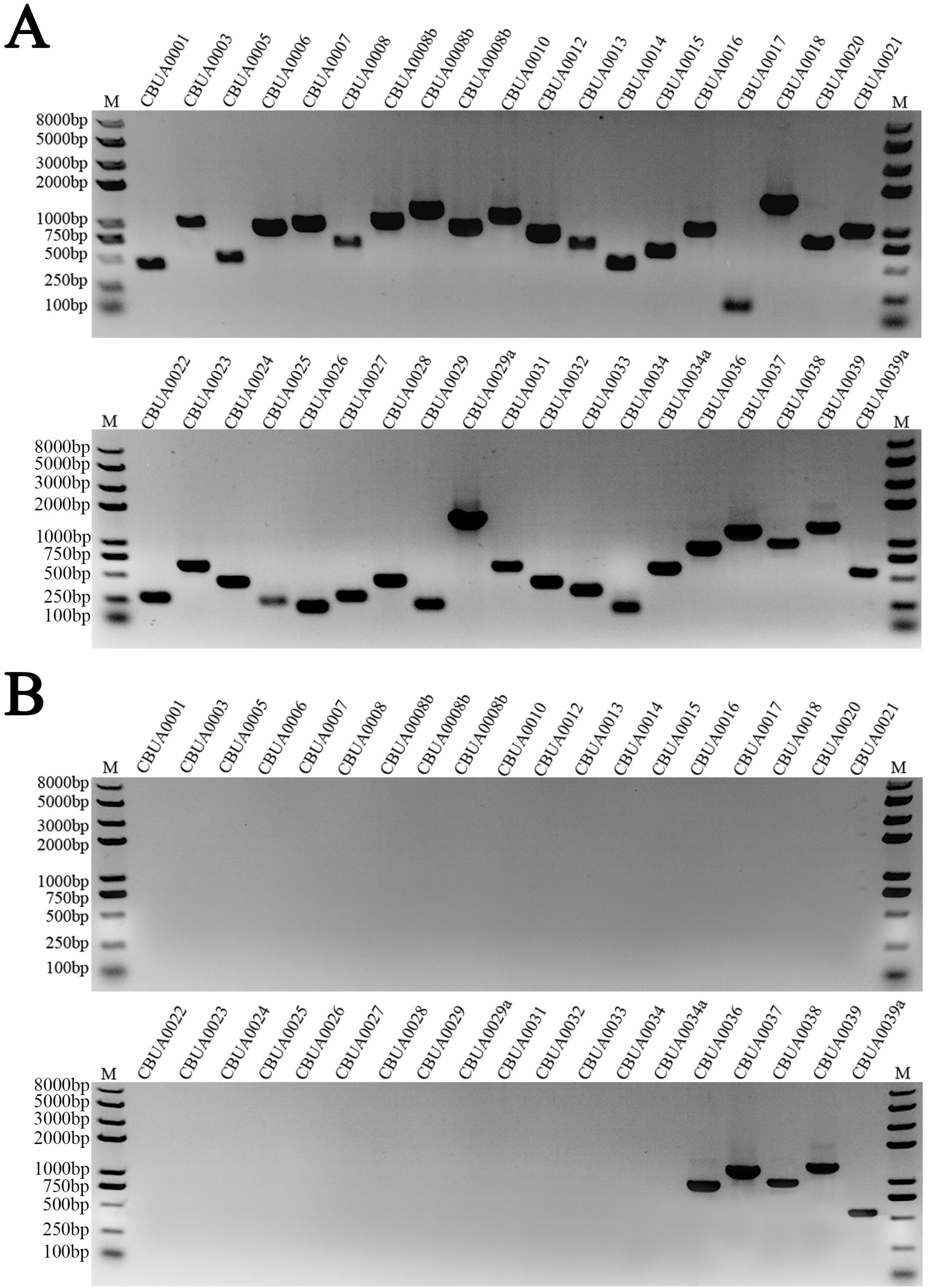
PCR detection of QpH1 ORFs in two transformants -NMII*pMMGK* (A) and NMII*pQGK* (B). All QpH1 ORFs are present in NMII*pMMGK* while only the CBUA0036-0039a ORFs can be detected in NMII*pQGK*.

### QpH1-deficient *C. burnetii* has normal growth kinetics in axenic media and BGMK cells

The transformants of pMMGK and pQGK -NMII*pMMGK* and NMII*pQGK* derived from a same Nine Mile II clone. In addition to carrying different shuttle vectors, their major genetic difference is the presence or absence of the endogenous QpH1 plasmid. Both shuttle vectors carry the eGFP cassette, and their clonal homogeneity was inspected by observing eGFP expression. We investigated the potential impact of QpH1 on bacterial growth with these two eGFP-labeled transformants. Their growth kinetics in ACCM-2 media and BGMK cells were assessed by measuring their one-step growth curves (Figure 4A-B). These two transformants display almost identical growth kinetics, suggesting QpH1 is not important for *C. burnetii* growth in axenic media and BGMK cells.

**Figure 4.**
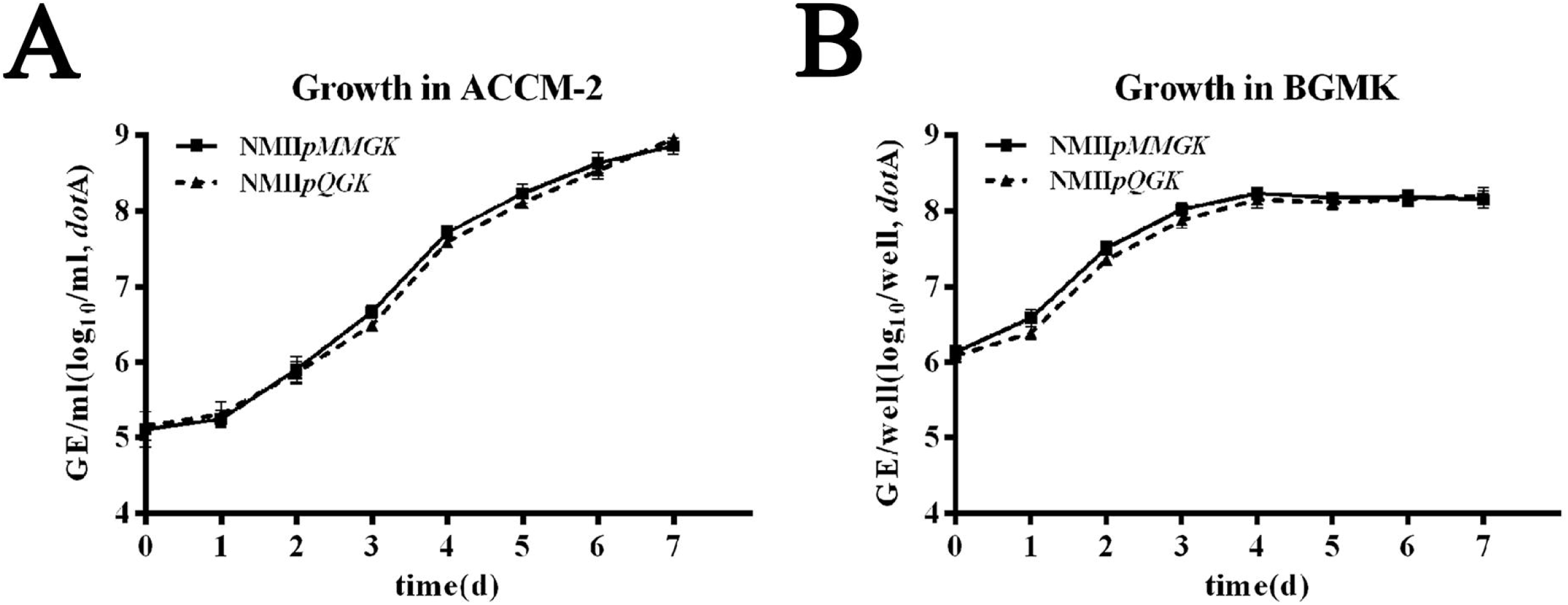
One-step growth curves of two transformants -NMII*pMMGK* (solid line) and NMII*pQGK* (dotted line) in axenic media (A) and BGMK cells (B). Growth was quantified by measuring GE. Results of axenic and cell cultures are from six and four separate replicates, respectively.

Interestingly, though having similar growth kinetics, these two transformants differ significantly in their expression of eGFP genes (Figure 5). The eGFP genes of these two transformants have identical promoters, but the fluorescence intensity of NMII*pMMGK* appears to be higher than that of NMII*pQGK*. The difference in level of eGFP expression between these two transformants may reflex the difference in gene copy numbers. The copy number of RSF1010-ori based shuttle vector is 3 to 6, while the QpH1 copy number is between one to three. Our data show that QpH1 and pQGK have a similar copy number, suggesting that pQGK retains all genes for QpH1 replication and partitioning. The copy number of pMMGK is about two to three times of the copy number of pQGK, which is consistent with their expression levels of eGFP. Overall, our data suggest the copy number of QpH1 and pQGK is under highly stringent control.

**Figure 5.**
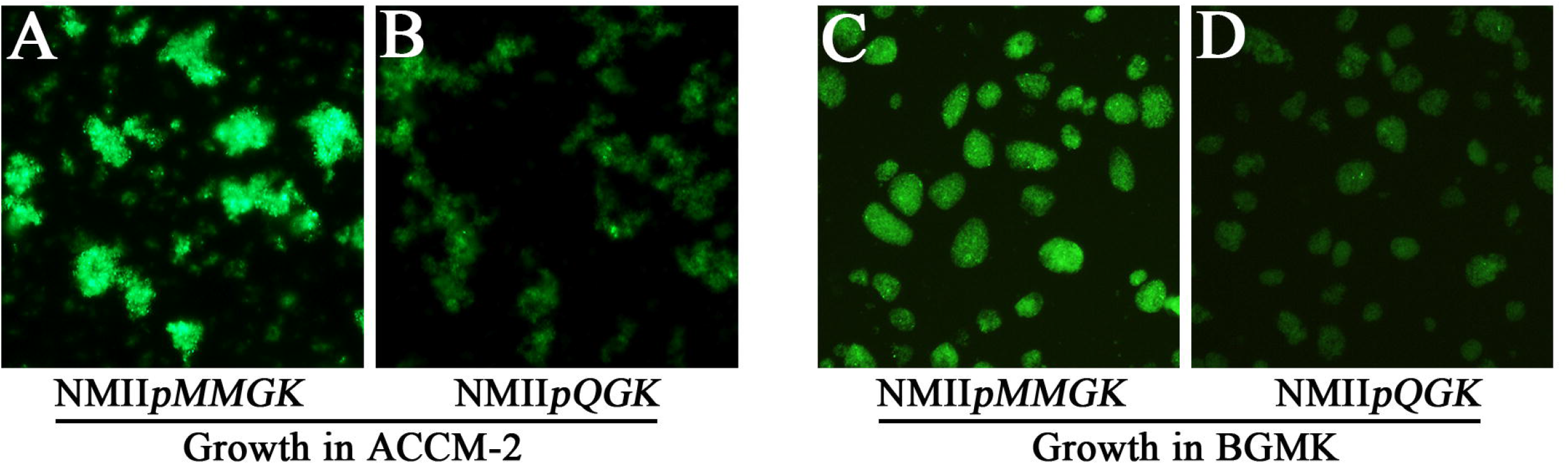
Phase and fluorescence microscopy of NMII*pMMGK* (A, C) and NMII*pQGK* (B, D) in axenic media (A-B) and BGMK cells (C-D). Compared to NMII*pQGK*, NMII*pMMGK* appears to have higher level of eGFP expression in both axenic media and BGMK cells.

### QpH1-deficient *C. burnetii* almost completely fails to colonize murine bone marrow derived macrophages

In vitro, *C. burnetii* can infect a variety of cell types including epithelial and fibroblast cell lines. However, this bacterium has an infection tropism for mononuclear phagocytes in natural infections [36]. We next characterized the growth of NMII*pMMGK* and NMII*pQGK* in human macrophage-like THP-1 cells and murine bone marrow-derived macrophages (murine BMDM) (Figure 6), both of which can support robust growth of *C. burnetii* phase II under normoxic culture conditions [37–39]. Equal MOIs of NMII*pMMGK* and NMII*pQGK* were used to infect THP-1 and murine BMDM cells.

**Figure 6.**
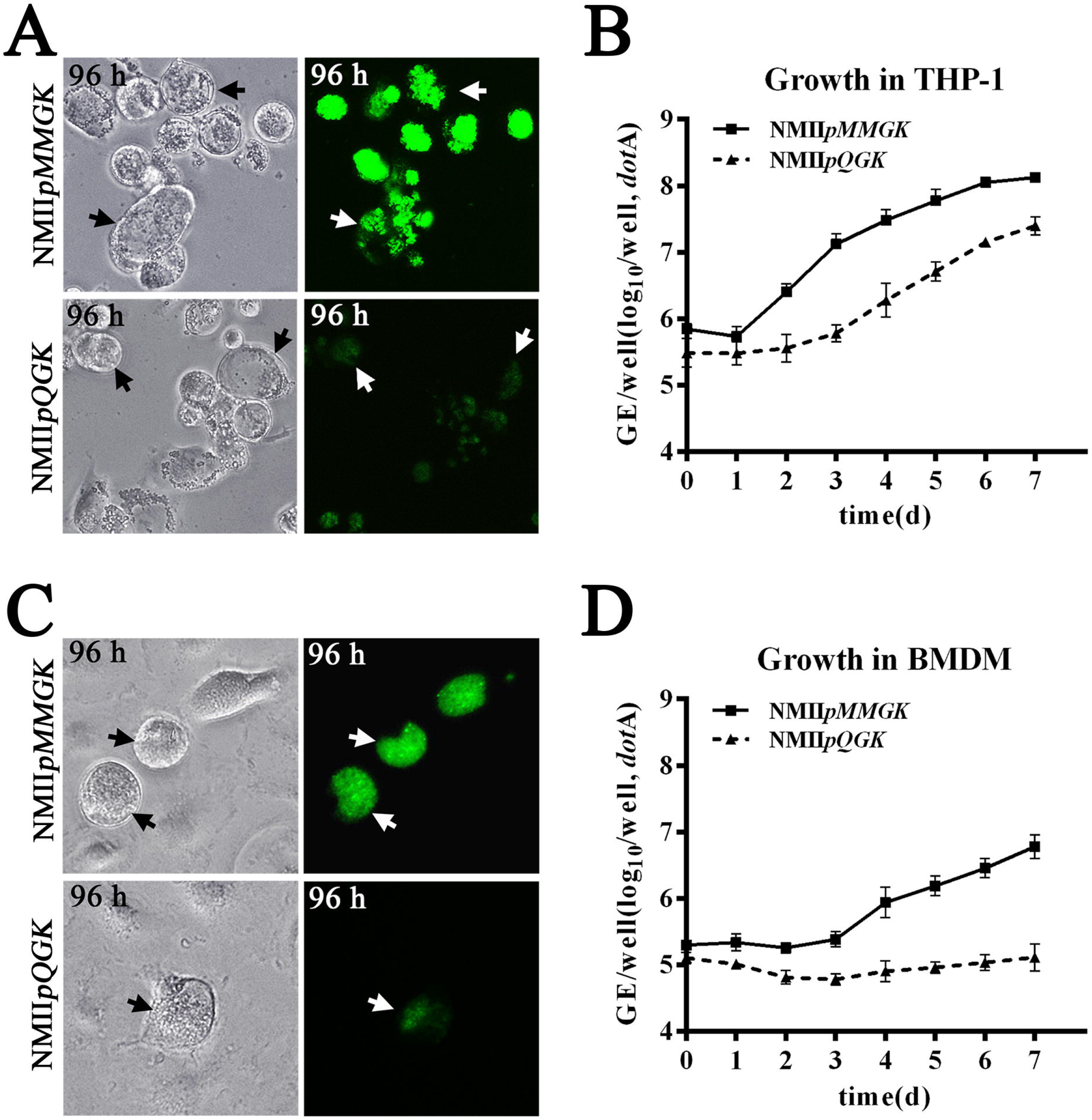
Growth of *C. burnetii* transformants NMII*pMMGK* and NMII*pQGK* on macrophage-like THP-1 cells (A, B) and bone marrow-derived murine macrophages (BMDMs) (C, D). (A) In THP-1 cells at 96 hours postinfection, CCVs of NMII*pMMGK* have strong eGFP expressions and are filled with bacterial particles while CCVs of NMII*pQGK* have weak eGFP expression and have much less bacterial particles. (B) Compared to NMII*pMMGK*, NMII*pQGK* has lower GEs at the initial attachment and lower growth yields throughout the culture period. (C) In BMDMs, NMII*pMMGK* forms normal-sized CCVs and has strong eGFP expression while NMII*pQGK* almost completely fails to colonize. A rarely observed CCV of NMII*pQGK* is shown. (D) In BMDMs, the GEs of NMII*pMMGK* increased close to two logs while the GEs of NMII*pQGK* have no significant growth throughout the culture period.

In THP-1 cells (Figure 6A), NMII*pMMGK* has robust growth at 96 hours post-infection, as shown by the densely packed bacterial clumps and intensive eGFP expressions in its large CCVs. In contrast, NMII*pQGK* CCVs are partially filled with bacterial clumps and have lower eGFP expression. This evident lower growth rate of NMII*pQGK* is confirmed in the one-step growth curves of these two strains (Figure 6B). It is noted that the lower eGFP expression of NMII*pQGK* in THP-1 is likely due to a combination of lower plasmid copy number and lower growth rate. In addition, we repeatedly measured a minor lower (p<0.05) number of NMII*pQGK* at the infection time point of zero, which is the starting time after bacterial entry (Figure 6B). Overall, these THP-1 infection results suggest that QpH1 plays roles in *C. burnetii* attachment and multiplication in human macrophages.

We next compared the growth of NMII*pMMGK* and NMII*pQGK* in murine BMDMs. In murine BMDMs at 96 hours post-infection, NMII*pMMGK* forms normal CCVs that can be easily observed in any microscope field (Figure 6C). This observation is consistent with previous finding of robust growth of *C. burnetii* phase II in murine BMDMs [38]. Contrarily, NMII*pQGK* produces almost zero CCVs. In our repeated experiments, only one NMII*pQGK* CCV was ever observed. This rarely observed CCV of NMII*pQGK* is shown in Figure 6C. Consistent with microscope observations, no growth yield of NMII*pQGK* is detected by qPCR (Figure 6D). It is noted our growth yields of NMII*pMMGK* are not optimal when compared to the report of Cockrell et al [38], which is likely due to our modified infection procedures, such as centrifugation at room temperature instead of 37°C. In addition, comparison of the GE numbers at the starting time point of infection suggested that NMII*pQGK* also has slightly reduced attachment efficiency (p<0.05), which is similar with findings in THP-1. Altogether, the murine BMDM infection results suggest that QpH1 is essential for *C. burnetii* survival in murine macrophages.

### Plasmid-less *C. burnetii* can be isolated by using CBUA0037 or -0038-deleted shuttle vectors

The shuttle plasmid pQGK can be stably maintained and has a normal plasmid copy number in *C. burnetii* Nine Mile phase II. We next investigated the plasmid genes for plasmid maintenance in pQGK. A PCR-based deletion mutagenesis approach was adopted to generate specific deletions in each of the five QpH1 ORFs (CBUA0036-0039, -0039a) (Figure 7A) [40]. The deletion regions range from 60% to 100% of these ORFs. Primers with unique restriction enzyme sites at their 5’ ends were used for PCR amplification as described in Materials and Methods. After digestion with restriction enzymes, purified PCR products were ligated and transformed into *E. coli* Trans5α. The desired plasmid construct was confirmed by sequencing. The plasmid knockout vectors are referred to as pQGK-D1, -D2, -D3, D4 and -D5, corresponding to deletion mutants of CBUA0036, -0037, -0038, -0039 and -0039a, respectively.

**Figure 7.**
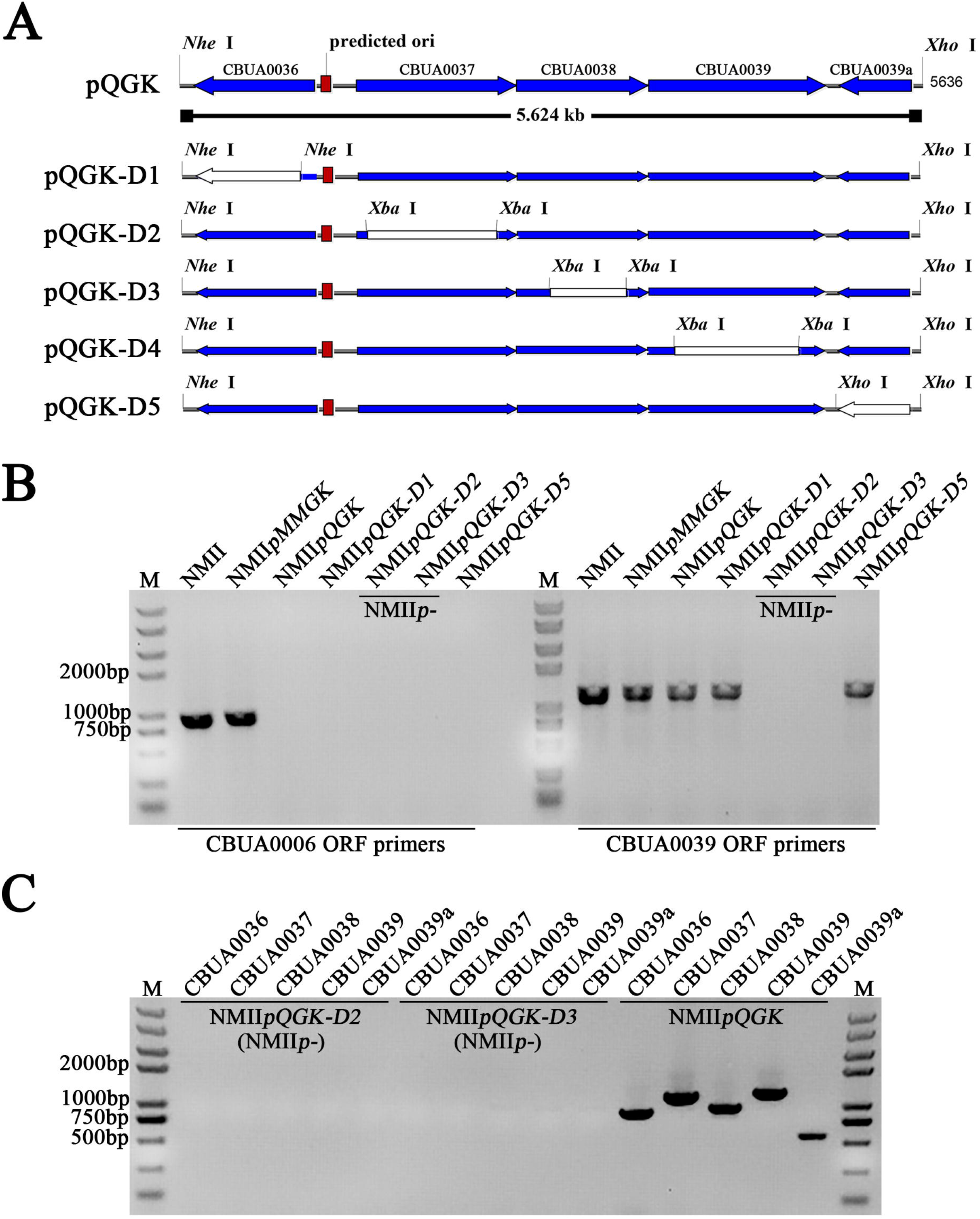
Construction of five individual QpH1 ORF knockout vectors and PCR identification of their transformants. (A) Schematic representation of deleted regions (white) of pQGK in five individual QpH1 ORF knockout vectors. The coding sequences of CBUA0036-0039a and their direction of transcription are shown by the arrows on the line. (B) The CBUA0006 ORF is absent in transformants of pQGK, -D1, -D2, -D3, -D5, suggesting these transformants are QpH1-deficient. The CBUA0039 ORF is also absent in transformants of pQGK-D2 and -D3, suggesting these two transformants contain no any plasmid. (C) The CBUA0036-0039a ORFs are absent in NMII*pQGK-D2* and NMII*pQGK-D3*, but present in NMII*pQGK*.

Transformations of *C. burnetii* Nine Mile phase II with these knockout vectors were performed. All knockout vectors contain the eGFP cassette. Their transformants were cultured in kanamycin media and were constantly observed by using fluorescence microscopy. After several passages, transformants were cloned for three times on semi-solid plates. The cloned transformants were then cultured in kanamycin media. The stability of the transformants was assessed by observing eGFP expression. Transformation results are summarized in Table 1. Stable transformants carrying pQGK-D1 and -D5 were isolated (Figure 7B). PCR detection of QpH1 genes indicate that QpH1 was also cured from these transformants. The results of pQGK-D1 and -D5 reveal that CBUA0036 and -0039a are not required for stable plasmid maintenance in axenic culture of *C. burnetii*.

**Table 1.** Viability and stability of *C. burnetii* Nine Mile phase II transformed with five pQGK deletion derivatives

| Plasmid | Phenotypes of transformants growing in kanamycin media |  |
| --- | --- | --- |
|  | Bacterial growth | eGFP stability |
| pQGK-D1<br>(ΔCBUA0036) | + | stable |
| pQGK-D2<br>(ΔCBUA0037) | + | unstable |
| pQGK-D3<br>(ΔCBUA0038) | + | unstable |
| pQGK-D4<br>(ΔCBUA0039) | - | transient |
| pQGK-D5<br>(ΔCBUA0039a) | + | stable |

Transformation with other three deletion vectors yielded transient or unstable transformants in the presence of antibiotic selection. For the transformation with pQGK-D4, green fluorescence was observed at the first passage after electroporation, but failed to expand in the presence of kanamycin selection. The green fluorescence can be repeatedly observed at the first passages in repeated experiments. This transient eGFP expression suggests that pQGK-D4 was introduced into *C. burnetii* by electroporation but cannot be stably maintained.

Transformation with pQGK-D2 and -D3 produced significant bacterial yields in the presence of kanamycin selection. However, clones from kanamycin plates always give a mixture of eGFP-positive and eGFP-negative bacterial clumps. Further identification of these eGFP negative clones revealed that these clones are double deficient of QpH1 and the deletion mutant of pQGK (Figure 7B-C). This unstable eGFP expression and the obtaining of plasmid-deficient *C. burnetii* suggest that pQGK-D2 and -D3 are incompatible with QpH1 and their transformants are constantly losing their shuttle plasmid due to partitioning deficiency. A plasmid-deficient *C. burnetii* clone NMII*p*-was isolated by three successive cloning of pQGK-D2 transformants. Altogether, these results show that CBUA0037, -0038 and -0039 are essential genes for QpH1 maintenance and the pQGK-D2 and -D3 vectors are valuable tools for constructing plasmid-deficient *C. burnetii*.

### QpH1-deficient *C. burnetii* has reduced pathogenicity in SCID mice

The SCID mouse model was established for identifying virulence factors in *C. burnetii* phase II [41]. We next analyzed the plasmid’s potential role in *C. burnetii* pathogenesis by using this model (Figure 8). Four *C. burnetii* strains -the parent Nine Mile phase II (NMII) and its three derivatives (NMII*pMMGK*, NMII*pQGK* and NMII*p*-) were used to infect SCID mice by peritoneal (IP) injections. Compared to PBS control group, all four *C. burnetii* strains induced different levels of splenomegaly in infected mice (Figure 8A-B). However, it is apparent that QpH1-bearing strains (NMII and NMII*pMMGK*) induced bigger splenomegaly that QpH1-deficient strains (NMII*pQGK* and NMII*p*-) (*p*<0.05), suggesting QpH1 plays a role in pathogenicity. It is noted that NMII*pMMGK* has a tendency of causing bigger splenomegaly than its parent NMII (*p*=0.10), indicative of enhanced virulence by the pMMGK plasmid. This enhanced virulence of NMII*pMMGK* is likely associated with this strain’s ability of normoxic growth. Characterization of *C. burnetii* normoxic growth was described in a separate manuscript [42]. Moreover, the sizes of splenomegaly are in accordance with *C. burnetii* genome equivalents in all infected mice (Figure 8C). It is also obvious that NMII*pQGK* retains partial pathogenicity in SCID mice, which is inexplicable as this strain has lost the ability of infecting mouse BMDMs generated from C57BL/6J. Altogether, our data suggest QpH1 plays a partial role in pathogenicity in the SCID mouse model.

**Figure 8.**
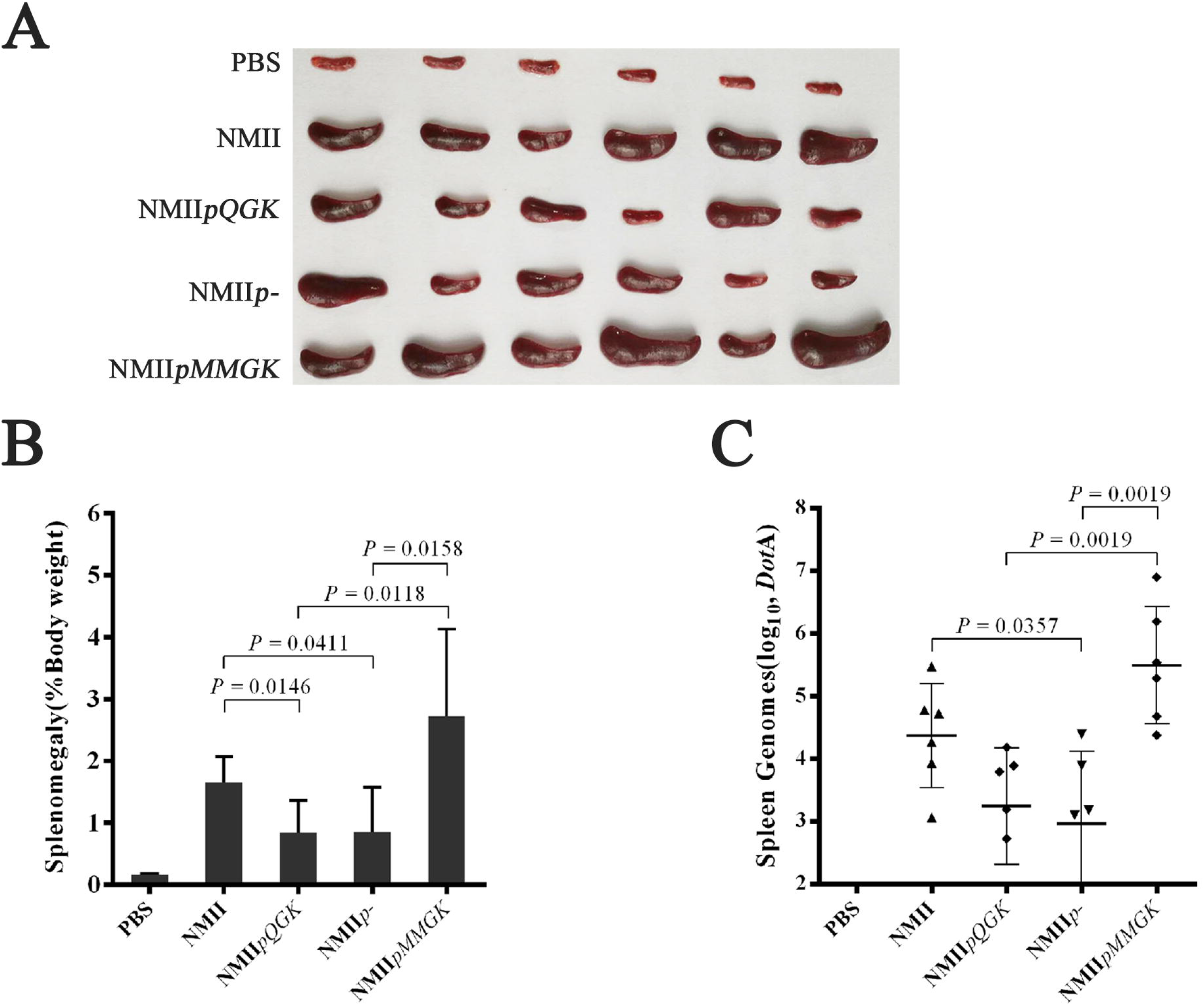
Splenomegaly of SCID mouse livers after intra-peritoneal challenge with *C. burnetii*. SCID mice were challenged with PBS or 1×10^7^ GE of *C. burnetii* via intra-peritoneal route and sacrificed 18 days after challenge. (A) Spleens were removed from control and challenged mice. (B) Splenomegaly calculated as spleen weight as a percentage of total body weight at the time of necropsy. (C) Genome equivalents calculated using TaqMan real-time PCR with DNA purified from infected spleens from 6 mice on 18 days after challenge with *C. burnetii*. For all panels, error bars represent standard deviations from the mean.

## Discussion

All *C. burnetii* has one of the four different types of plasmids or one plasmid-like chromosomally integrated sequence. The plasmids’ role in *C. burnetii* biology has been implicated by the identification of type IV secretion effectors [4, 19]. Our goal was to construct and characterize plasmid-deficient *C. burnetii*. We show that QpH1-deficient *C. burnetii* Nine Mile phase II -NMII*pQGK* can be constructed by transformation with a QpH1-based shuttle vector pQGK. The NMII*pQGK* strain has normal growth in axenic media and BGMK cells, displays partial growth defects in human macrophages and, most strikingly, shows severe infection-deficiency in mouse BMDMs. In the SCID mouse model, the NMII*pQGK* strain has partially reduced pathogenicity. We show that CBUA0037, -0038 and -0039 are essential ORFs for QpH1 maintenance, and unmarked QpH1-deficient mutants can be made by transformation with an unstable CBUA0037, or CBUA0038 deletion shuttle vector.

Plasmids are commonly classified by incompatibility (Inc) typing [43]. Plasmids that share the same replication mechanisms are incompatible. The Inc type of *C. burnetii* plasmids remains to be determined. An incompatible plasmid can be used for making plasmid-deficient *C. burnetii* strains. Our findings that pQGK can cure the endogenous QpH1 by transformation and has a same copy number of QpH1 indicate that pQGK retains an intact QpH1 replicon including genes for plasmid replication, partitioning and copy number control. The pQGK plasmid can be used for Inc typing of *C. burnetii* plasmids. The pQGK derivatives -pQGK-D2 or -D3 can be used for making unmarked plasmid-deficient mutants. *C. burnetii* has four types of plasmids. This plasmid curing approach has potential applications for investigating plasmid-type virulence of *C. burnetii*.

Plasmids can maintain their copy numbers by one of the three general classes of negative regulatory systems [44]. Iterons and their cognate replication (Rep) initiator proteins comprise the iteron-involved negative control system [45]. Our results revealed that QpH1 is of low copy number and has atypical iterons. The replication and copy number control of QpH1 is likely regulated through the iterons-Rep complex. Iterons are able to exert plasmid incompatibility [46]. Further studies are required to understand the iterons involved regulation of plasmid replication in *C. burnetii*. In addition, these iterons’ cognate RepA protein -annotated CBUA0039, is supported by our finding of transient transformation with the pQGK-D4 vector. An AT-rich region, locating between the iterons and CBUA0037, could be another essential element for QpH1 replication [47].

QpH1 is a plasmid of low copy number [10]. Low-copy plasmids have an active partitioning system. Two putative partitioning proteins -CBUA0036 (ParB) and -0037 (ParA), have been annotated based on protein homology. Our results suggest that CBUA0037 and -0038 are essential partitioning proteins while CBUA0036 is nonessential for plasmid maintenance. The commonly used RSF1010-ori based shuttle vectors for *C. burnetii* is also used in the genetic studies of *Legionella pneumophila* [33]. Our identification of essential genes for QpH1 maintenance might be useful for constructing QpH1 based shuttle vectors that can be stably maintained in *L. pneumophila*.

Besides genes for plasmid maintenance (replication, copy number control and partitioning), plasmids from pathogenic microbes often contain genes associated with virulence and/or antibiotic resistance [40]. Three of the 40 QpH1 ORFs are essential for plasmid maintenance. Most of other ORFs likely contribute to *C. burnetii* virulence in various ways. Our finding of QpH1-deficient *C. burnetii* failing to colonize murine BMDMs likely explains the high conservation of plasmid genes in *C. burnetii*. We postulate that *C. burnetii* plasmids are essential for bacterial survival in natural reservoir hosts, including rodents. The role of rodents as a potential reservoir was suggested by finding *C. burnetii* infections in mice in different areas and countries [48–50].

How might plasmid genes contribute to the colonization of murine BMDMs? Previous studies using human cells found that the type IV secretion system and a repertoire of type IV effectors required for the CCV biogenesis and *C. burnetii* growth [25–28]. The role that these type IV effectors play in *C. burnetii* pathogenicity is of intense interest [51]. Plasmid encoded effectors are possibly involved in the reduced growth efficiency of QpH1-deficient *C. burnetii* on THP-1 cells, and might also relate to the failed colonization of murine BMDMs by QpH1-deficient *C. burnetii*. Newton and co-workers found that effector proteins can be translocated as early as 1 h post-infection in murine BMDMs, while effector translocation can only be detected after 8 h post-infection in HeLa cells [52]. We surmise that early translocation of effectors might play an essential role in *C. burnetii* resisting antimicrobial activities of murine BMDMs. Further studies to investigate the pathogenesis of plasmid genes required for *C. burnetii* infection and growth in murine BMDMs are warranted.

The QpH1-deficient *C. burnetii* almost completely abrogates its capacity of infecting murine BMDMs. Thus, it is unexpected that this mutant has only partially reduced its pathogenicity in SCID mice. It is possible that the antimicrobial capacity of macrophages is also dampened in this immunodeficiency mouse model. The immunocompetent animal model with *C. burnetii* phase I as control may be more appropriate for studying plasmid pathogenesis [12]. It is also unexpected that the pMMGK transformant appears to have enhanced pathogenicity. Though similar with other RSF1010-ori based shuttle vectors, the pMMGK vector confers an unexpected capability of normoxic growth on *C. burnetii*, which likely relates to the transformant’s enhanced virulence. The normoxic growth of *C. burnetii* was characterized in a separate manuscript [42].

In conclusion, we investigated the functions of *C. burnetii* QpH1 plasmid genes. We report that QpH1 is a low copy plasmid. We propose that the intergenic region between CBUA0036 and -0037 is the plasmid’s origin of replication, and the plasmid’s copy number is regulated through atypical iterons within this region. The CBUA0037, -0038 and -0039 ORFs encode two partitioning proteins and Rep proteins, respectively. QpH1 can be cured by transformation with pQGK or its two derivatives (pQGK-D2 and -D3). We present evidence that QpH1 is nonessential for *C. burnetii* growth in axenic media and BGMK cells. QpH1-deficient *C. burnetii* phase II has minor growth deficiency in human macrophages. Most importantly, we found QpH1 is essential for *C. burnetii* colonization of murine BMDMs, highlighting an important role of the plasmid for *C. burnetii* persistence in rodents. It is certain that more plasmid related phenotypes remain to be identified. Our shuttle vectors will further the progress of genetic studies of *C. burnetii* and its plasmids. This work represents an important step toward unravelling the mystery of plasmid conservation in *C. burnetii*.

## Methods

### Bacterial strains, cell lines and key reagents

*C. burnetii* Nine Mile phase II (NM II) is from our laboratory strain collection and was passaged in ACCM-2 as previously described [22]. *E. coli* Trans5α (TransGen Biotech) was used for recombinant DNA procedures and was grown in Luria-Bertan (LB) media. Buffalo green monkey kidney cells (BGMK) were grown in RPMI 1640 medium supplemented with 10% FBS. Human monocytic leukemia cells (THP-1) were cultured in high-glucose-containing Dulbecco’s Modified Eagle Medium (DMEM) supplemented with 10% FBS. Murine bone marrow-derived macrophages (BMDM) were generated from C57BL/6J female mice as described by Cockrell et al [38]. Primers used in this study are listed in S1 Table. *Escherichia coli* Trans5α (TransGen Biotech) was used for all plasmid amplification.

### Construction of a RSF1010 ori-based shuttle vector pMMGK

RSF1010 ori-based shuttle vectors are commonly used for *C. burnetii* transformation [30, 33]. We constructed a RSF1010 ori-based vector pMMGK based on the backbone of pMMB207 (kindly provided by Xuehong Zhang from Shanghai Jiaotong University, Shanghai, China). Promoter regions of CBU0311 and CBU1169 were amplified with primer pairs P311-F/P311-R and P1169-F/P1169-R, respectively. The Kan^R^ and eGFP genes were amplified from pEASY-T1 and pEGFP-C1, with primer pairs Kan-F/Kan-R and eGFP-F/eGFP-R, respectively. Overlapping PCR with primers P311-F/Kan-R was used to produce the P311-eGFP-P1169-Kan^R^ segment. Two segments -pUC ori and ampicillin resistance gene (Amp^R^) were amplified from pEASY-T1 with primer pairs pUC-F/pUC-R and Amp-F/Amp-R, respectively. Overlapping PCR with primers P311-F/pUC-R was used to produce the P311-eGFP-P1169-Kan^R^-pUC ori segment. This segment and the Amp^R^-containing segment were digested with *Nhe* I and *Xho* I, and ligated to generate the vector pAGK. The RSF1010 backbone and the P311-eGFP-P1169-Kan^R^ segment were amplified from pMMB207 and pAGK, with primer pairs RSF1010-F/RSF1010-R and P311-F/KAN-R, respectively, then were digested with *Nhe* I and *Xho* I and ligated to generate the shuttle vector pMMGK (S2 Appendix).

### Construction of *C. burnetii* QpH1-based shuttle vector pQGK

The shuttle vector pQGK was constructed by replacing the Amp^R^ region of pAGK with a predicted region for plasmid replication and partitioning in QpH1. Briefly, a 5.6-kb segment covering CBUA0036-39a was amplified with primers pQ-F and pQ-R. PCR amplifications were performed in 100-μl reaction volumes containing 50 ng DNA, a 0.2 mM concentration of each dNTP, a 0.2 μM concentration of each primer, 2.5 U TransStart FastPfu DNA polymerase (TransGen Biotech), and 20 μl buffer. PCR conditions were as follows: initial denaturation at 95°C for 4 min followed by 2 cycles of amplification at 95°C for 30s, 55°C for 30s and 72°C for 1 min every two kb and then 28 cycles of amplification at 95°C 30s, 57°C for 30s and 72°C for 1 min every two kb and a final extension at 72°C for 10 min. This QpH1 segment and the pAGK plasmid were digested with *Nhe* I and *Xho* I and ligated to produce the shuttle plasmid pQGK (S3 Appendix).

### Construction of deletion derivatives of pQGK

Deletion derivative of pQGK were constructed using a modified PCR-based approach [40]. Primers pairs for knocking out CBUA0036-39a were shown in Table S1 in the supplemental material. Briefly, PCR amplifications were performed using TransStart FastPfu DNA polymerase (TransGen Biotech) and PCR conditions were described as above. PCR products were purified by a QIAquick PCR purification kit (Qiagen) and doubly digested with DMT (to remove template DNA, TransGen Biotech) and restriction enzymes as indicated. Digested PCR products were ligated using Quick T4 DNA ligase (NEB) and transformed into *E. coli* Trans5α. Clones were screened for plasmids containing QpH1 ORFs but lacking the deleted region by PCR.

### Transformation of *C. burnetii* Nine Mile phase II

All plasmids used in this study were purified using a plasmid Maxi Kit (Qiagen) and subjected to sequencing confirmation. The resulting vector was transformed into *C. burnetii* Nine Mile phase II by electroporation [31]. For every passage after electroporation, *C. burnetii* was cultured at 2.5% O2 and 5% CO_2_ for seven days in ACCM-2 containing 400 μg/mL kanamycin. Fluorescence microscopy was conducted to check eGFP expression after each passage. When eGFP expression was easily observed in bacterial clumps, semi-solid agarose plates were then used for three successive cloning. *C. burnetii* transformants were expanded in ACCM-2, suspended in SPG (0.25M sucrose, 10mM sodium phosphate, 5mM L-glutamic acid) and kept frozen at −80°C. Genome copy numbers were determined by Taqman probe qPCR specific to *dotA* [53].

### PCR identification of transformants

The presence or absence of QpH1 in *C. burnetii* transformants were detected by PCR. Total DNAs were extracted using DNeasy blood & tissue kit (Qiagen), then used as templates for PCR detection with TransTaq high fidelity DNA polymerase (TransGen) and QpH1 gene primers (S4 Table). PCR conditions are as follows: initial denaturation at 94°C for 5 min, followed by 35 cycles of 94°C for 30 s, 55°C for 30 s, 72°C for 90 s.

### One-step growth curves

To measure the one-step growth curves in axenic media, seed stocks of *C. burnetii* was inoculated at a density of 2×10^6^ GE/ml in 96-well plates containing 150 μl ACCM-2 medium per well. Cultures were incubated in a CO-170 tri-gas incubator at 37°C with 2.5% O2 and 5% CO_2_. Samples of axenic culture were collected at indicated time points, then were centrifuged at 20,000 *g* for 10 min to harvest bacteria.

To measure the one-step growth curves in cells, seed stocks of *C. burnetii* was plated onto 24-well cell (BGMK, THP-1 and BMDM) monolayers at an MOI of 50. Then, 24-well plates were centrifuged at 800 *g* for 25 min at room temperature, placed in a CO_2_ incubator (5% CO_2_) at 37°C for 2 hours. Cells were then washed twice with PBS and 1 ml cell medium was added to each well. At indicated time, cell samples were harvested by using trypsin (Hyclone) treatment and centrifugation.

Total DNAs were extracted with DNeasy Blood Tissue Kit (Qiagen). Genome copy numbers of *C. burnetii* in both axenic and cell cultures were determined by Taqman probe qPCR as previously described [53]. PCR conditions were as follows: initial denaturation at 94°C for 10min, followed by 40 cycles of amplification at 94°C for 15 s, 60°C for 1min. All samples were performed in quadruplicate.

### SCID mice infections

SCID (CB17/Icr-*Prkdc^scid^*/IcrlcoCrl) mice were purchased from Vital River Laboratory Animal Technology Co., Ltd (Beijing, China). Freshly cultured *C. burnetii* and its transformants –NMII*p*-, NMII*pMMGK* and NMII*pQGK* were diluted to 5×10^7^ GE/mL in PBS. Six six-week old female mice per group were infected with 1×10^7^ GE *C. burnetii* in 200 μl PBS via intra-peritoneal injection. At 18 days postinfection, hearts, lungs, livers, and spleens were aseptically harvested and stored at −80°C, additionally, spleens were weighted to determine splenomegaly (spleen weight/body weight). All samples were individually added up to 2 mL with PBS and homogenized using a glass grinder. DNAs were purified from 20 μL tissue homogenate with DNeasy Blood Tissue Kit (Qiagen) and were used to determine *C. burnetii* GE by qPCR as described above. All animal experimental procedures involved were performed according to protocols approved by the Institutional Animal Care and Use Committee of Fifth Medical Center, General Hospital of Chinese PLA. The project ethics review approval number is IACUC-2018-0020.

## Acknowledgments

This work was supported by National Natural Science Foundation of China (No. 31570177), Fundamental Research Funds for Central Universities (No. BUCTRC201917), Key Project of Beijing University of Chemical Technology (No. XK1803-06) and Foundation of State Key Laboratory of Pathogen and Biosecurity (No. SKLPBS1409).

## Supporting information

**S1 Table. Primers for plasmids construction.**

**S2 Appendix. Complete sequence of the pMMGK plasmid.**

**S3 Appendix. Complete sequence of the pQGK plasmid**

**S4 Table. Primers for PCR Identification.**

