## Supplementary material for "The *Coxiella burnetii* QpH1 plasmid is a virulence factor for colonizing bone marrow-derived murine macrophages": S1 Table.docx

**S1 Table. Primers for plasmids construction**

| **Primers** | **Sequence (5’ to 3’) (restriction sites in red)** |
| --- | --- |
| For shuttle vector construction | |
| P311-F  P311-R | CCAATGTTGCTAGCGATTATTAATTCAAACGGGTC  CTCGCCCTTGCTCACCATGTCAAATCTCCGTTTTCAAC |
| GFP-F  GFP-R | GTTGAAAACGGAGATTTGACATGGTGAGCAAGGGCGAG  GAACCTGTTTGGATCCTTACTTGTACAGCTCGTCCATG |
| P1169-F  P1169-R | CTGTACAAGTAAGGATCCAAACAGGTTCTCTAATTAATC  CAATCATATGCGCTCTCCTTTCAG |
| Kan-F  Kan-R  KAN-R | TCTGAAAGGAGAGCGCATATGATTGAACAAGATGGAT  GACAGGGTACCTTATTAGAAGAACTCGTCAAGAAG  TTCCTACCTCGAGTTAGAAGAACTCGTCAAGAAG |
| Amp-F  Amp-R | ACGTCGCTCTCGAGTTATTACCAATGCTTAATCAGTG  CTAGTTAGTCGCTAGCTATTTTCTCCTTACGCATCTG |
| pUC-F  pUC-R | CTTCTAATAAGGTACCCTGTCAGACCAAGTTTACTCAT  CATACCTCAGCTCGAGGGGATAACGCAGGAAAGAAC |
| RSF1010-F  RSF1010-R | GACACGACGCACCTCGAGAGTTATTGTCTTCAAATTCCCGT  TATCTCCGCGTGCTAGCAGCTGTGCGGCAGCGCTCAGTAG |
| pQ-F  pQ-R | GCATTGCTGCTAGCCAACCTCCCTTCTTTCCCAATC  CTTCTACATCTCGAGATTCTTTGTAGAATCGCTGTC |
| For ORF deletion derivatives construction | |
| pQ36-NheIF  pQ36-NheIR | GCTTCATCAAGCTAGCAGAGGCGTTCGCTAATGACTTC  CCAATGTTGCTAGCGATTATTAATTCAAACGGGTC |
| pQ37-XbaIF  pQ37-XbaIR | TTCTATGTGACCTCTAGATCTATGATGTATCAAAACCACGAG  CTGGACTCGTCGTCTAGATTACCAGCTTGGTAGAATTTGTC |
| pQ38-XbaIF  pQ38-XbaIR | AATCTATCGATCTCTAGATAAGACTAAAGATGGGAAAAAGC  TAGCTCGATCGCTCTAGATTTATATTGCCAAGTTCCTCATC |
| pQ39-XbaIF  pQ39-XbaIR | CTGAAGCTCTCTTCTAGACTAGTGATGGATTTTGAGGTTG  TGTGCACATCTCTCTAGATTGTAGGGTAACTCTGGATAAG |
| pQ39a-XhoIF  pQ39a-XhoIR | CATACCTCAGCTCGAGGGGATAACGCAGGAAAGAAC  TACGAATATCTCGAGAAAGTCTCCAAAGAGAACATCCTG |
