## Supplementary material for "The *Coxiella burnetii* QpH1 plasmid is a virulence factor for colonizing bone marrow-derived murine macrophages": S2 Appendix.DOCX

Complete sequence of the pMMGK plasmid:

CTCGAGTTAGAAGAACTCGTCAAGAAGGCGATAGAAGGCGATGCGCTGCGAATCGGGAGCGGCGATACCGTAAAGCACGAGGAAGCGGTCAGCCCATTCGCCGCCAAGCTCTTCAGCAATATCACGGGTAGCCAACGCTATGTCCTGATAGCGGTCCGCCACACCCAGCCGGCCACAGTCGATGAATCCAGAAAAGCGGCCATTTTCCACCATGATATTCGGCAAGCAGGCATCGCCATGGGTCACGACGAGATCCTCGCCGTCGGGCATGCGCGCCTTGAGCCTGGCGAACAGTTCGGCTGGCGCGAGCCCCTGATGCTCTTCGTCCAGATCATCCTGATCGACAAGACCGGCTTCCATCCGAGTACGTGCTCGCTCGATGCGATGTTTCGCTTGGTGGTCGAATGGGCAGGTAGCCGGATCAAGCGTATGCAGCCGCCGCATTGCATCAGCCATGATGGATACTTTCTCGGCAGGAGCAAGGTGGGATGACAGGAGATCCTGCCCCGGCACTTCGCCCAATAGCAGCCAGTCCCTTCCCGCTTCAGTGACAACGTCGAGCACAGCTGCGCAAGGAACGCCCGTCGTGGCCAGCCACGATAGCCGCGCTGCCTCGTCCTGCAGTTCATTCAGGGCACCGGACAGGTCGGTCTTGACAAAAAGAACCGGGCGCCCCTGCGCTGACAGCCGGAACACGGCGGCATCAGAGCAGCCGATTGTCTGTTGTGCCCAGTCATAGCCGAATAGCCTCTCCACCCAAGCGGCCGGAGAACCTGCGTGCAATCCATCTTGTTCAATCATATGCGCTCTCCTTTCAGAAGGATTAATGTCATTATTTATTTATGGGGTATGGAGAGGGATATTTCAAGGCGCACAAATATGATTCCAGGCTCGTTCTGCGGTTCCGCCTGGGTGTTCGCTGAGATAGGTTTTTACTTCGTTATACGAAAGCGGCGAATCGCCAAAAATTTTTTTTCCAAGTTCGCTGCGAAACGAAGCCATTTCCCGAAGAAGTATGTGCCAAGTGTTTGTAGGTTTTGTTAAAATTTTCTAATAAGGATAATAGTTCATTGCACTTGTCCGTCACAAAGGTTATTTTATCCACAAAAAATTTGCCTACTAGGCCCTCCATTTCAGGAGATAAGAACGCATCATTTATCCATTTGAGTTTTTGGAATGGTAAAGTTTTTAAAAAATCATCTATCTCTGAGGTTTCTTTAGATAGAAGCTTTGGTAGAAAAACATTGTCCGGATTAATTAGAGAACCTGTTTGGATCCTTACTTGTACAGCTCGTCCATGCCGAGAGTGATCCCGGCGGCGGTCACGAACTCCAGCAGGACCATGTGATCGCGCTTCTCGTTGGGGTCTTTGCTCAGGGCGGACTGGGTGCTCAGGTAGTGGTTGTCGGGCAGCAGCACGGGGCCGTCGCCGATGGGGGTGTTCTGCTGGTAGTGGTCGGCGAGCTGCACGCTGCCGTCCTCGATGTTGTGGCGGATCTTGAAGTTCACCTTGATGCCGTTCTTCTGCTTGTCGGCCATGATATAGACGTTGTGGCTGTTGTAGTTGTACTCCAGCTTGTGCCCCAGGATGTTGCCGTCCTCCTTGAAGTCGATGCCCTTCAGCTCGATGCGGTTCACCAGGGTGTCGCCCTCGAACTTCACCTCGGCGCGGGTCTTGTAGTTGCCGTCGTCCTTGAAGAAGATGGTGCGCTCCTGGACGTAGCCTTCGGGCATGGCGGACTTGAAGAAGTCGTGCTGCTTCATGTGGTCGGGGTAGCGGCTGAAGCACTGCACGCCGTAGGTCAGGGTGGTCACGAGGGTGGGCCAGGGCACGGGCAGCTTGCCGGTGGTGCAGATGAACTTCAGGGTCAGCTTGCCGTAGGTGGCATCGCCCTCGCCCTCGCCGGACACGCTGAACTTGTGGCCGTTTACGTCGCCGTCCAGCTCGACCAGGATGGGCACCACCCCGGTGAACAGCTCCTCGCCCTTGCTCACCATGTCAAATCTCCGTTTTCAACTAAAGTTAAAACAAAATCAAAATAAGCCTATATTTAAGCCTATCTTAGCTTTTCTCAAAGCTATAAAATGAAGAATATCGATATCTTTTTTAAATGTCAACACTTAAATCCCATAAAATCATTAAAATAAGAAAAATCTAAGCTTCTACCGATAATTAAGGATATTCAATTACAAAAAAAAAATGCCCATTAATTAAACAATCCGTTTAATTAATGGGCATTTGCTTTCTATTTTGTGTAATTAGGAATCTAATAATAATTCATTTAGCCCTTTCACAAAACTGGCAGGGTCTTTAAGCTGCTCTCCTTCAGCTAGGAGCGCTTGATTAAGCAATAAATCGGCCCAACGATTAAAGCGTGTTTTATCAGATTCATTTTTCACCCGTAAAATTAAGGGGTGCGATGGATTAATTTCTAAAATAGGTTTTGCCTGCATGAAATCCTGACCCGTTTGAATTAATAATCGCTAGCAGCTGTGCGGCAGCGCTCAGTAGGCAATTTTTCAAAATATTGTTAAGCCTTTTCTGAGCATGGTATTTTTCATGGTATTACCAATTAGCAGGAAAATAAGCCATTGAATATAAAAGATAAAAATGTCTTGTTTACAATAGAGTGGGGGGGGTCAGCCTGCCGCCTTGGGCCGGGTGATGTCGTACTTGCCCGCCGCGAACTCGGTTACCGTCCAGCCCAGCGCGACCAGCTCCGGCAACGCCTCGCGCACCCGCTTGCGGCGCTTGCGCATGGTCGAACCACTGGCCTCTGACGGCCAGACATAGCCGCACAAGGTATCTATGGAAGCCTTGCCGGTTTTGCCGGGGTCGATCCAGCCACACAGCCGCTGGTGCAGCAGGCGGGCGGTTTCGCTGTCCAGCGCCCGCACCTCGTCCATGCTGATGCGCACATGCTGGCCGCCACCCATGACGGCCTGCGCGATCAAGGGGTTCAGGGCCACGTACAGGCGCCCGTCCGCCTCGTCGCTGGCGTACTCCGACAGCAGCCGAAACCCCTGCCGCTTGCGGCCATTCTGGGCGATGATGGATACCTTCCAAAGGCGCTCGATGCAGTCCTGTATGTGCTTGAGCGCCCCACCACTATCGACCTCTGCCCCGATTTCCTTTGCCAGCGCCCGATAGCTACCTTTGACCACATGGCATTCAGCGGTGACGGCCTCCCACTTGGGTTCCAGGAACAGCCGGAGCTGCCGTCCGCCTTCGGTCTTGGGTTCCGGGCCAAGCACTAGGCCATTAGGCCCAGCCATGGCCACCAGCCCTTGCAGGATGCGCAGATCATCAGCGCCCAGCGGCTCCGGGCCGCTGAACTCGATCCGCTTGCCGTCGCCGTAGTCATACGTCACGTCCAGCTTGCTGCGCTTGCGCTCGCCCCGCTTGAGGGCACGGAACAGGCCGGGGGCCAGACAGTGCGCCGGGTCGTGCCGGACGTGGCTGAGGCTGTGCTTGTTCTTAGGCTTCACCACGGGGCACCCCCTTGCTCTTGCGCTGCCTCTCCAGCACGGCGGGCTTGAGCACCCCGCCGTCATGCCGCCTGAACCACCGATCAGCGAACGGTGCGCCATAGTTGGCCTTGCTCACACCGAAGCGGACGAAGAACCGGCGCTGGTCGTCGTCCACACCCCATTCCTCGGCCTCGGCGCTGGTCATGCTCGACAGGTAGGACTGCCAGCGGATGTTATCGACCAGTACCGAGCTGCCCCGGCTGGCCTGCTGCTGGTCGCCTGCGCCCATCATGGCCGCGCCCTTGCTGGCATGGTGCAGGAACACGATAGAGCACCCGGTATCGGCGGCGATGGCCTCCATGCGACCGATGACCTGGGCCATGGGGCCGCTGGCGTTTTCTTCCTCGATGTGGAACCGGCGCAGCGTGTCCAGCACCATCAGGCGGCGGCCCTCGGCGGCGCGCTTGAGGCCGTCGAACCACTCCGGGGCCATGATGTTGGGCAGGCTGCCGATCAGCGGCTGGATCAGCAGGCCGTCAGCCACGGCTTGCCGTTCCTCGGCGCTGAGGTGCGCCCCAAGGGCGTGCAGGCGGTGATGAATGGCGGTGGGCGGGTCTTCGGCGGGCAGGTAGATCACCGGGCCGGTGGGCAGTTCGCCCACCTCCAGCAGATCCGGCCCGCCTGCAATCTGTGCGGCCAGTTGCAGGGCCAGCATGGATTTACCGGCACCACCGGGCGACACCAGCGCCCCGACCGTACCGGCCACCATGTTGGGCAAAACGTAGTCCAGCGGTGGCGGCGCTGCTGCGAACGCCTCCAGAATATTGATAGGCTTATGGGTAGCCATTGATTGCCTCCTTTGCAGGCAGTTGGTGGTTAGGCGCTGGCGGGGTCACTACCCCCGCCCTGCGCCGCTCTGAGTTCTTCCAGGCACTCGCGCAGCGCCTCGTATTCGTCGTCGGTCAGCCAGAACTTGCGCTGACGCATCCCTTTGGCCTTCATGCGCTCGGCATATCGCGCTTGGCGTACAGCGTCAGGGCTGGCCAGCAGGTCGCCGGTCTGCTTGTCCTTTTGGTCTTTCATATCAGTCACCGAGAAACTTGCCGGGGCCGAAAGGCTTGTCTTCGCGGAACAAGGACAAGGTGCAGCCGTCAAGGTTAAGGCTGGCCATATCAGCGACTGAAAAGCGGCCAGCCTCGGCCTTGTTTGACGTATAACCAAAGCCACCGGGCAACCAATAGCCCTTGTCACTTTTGATCAGGTAGACCGACCCTGAAGCGCTTTTTTCGTATTCCATAAAACCCCCTTCTGTGCGTGAGTACTCATAGTATAACAGGCGTGAGTACCAACGCAAGCACTACATGCTGAAATCTGGCCCGCCCCTGTCCATGCCTCGCTGGCGGGGTGCCGGTGCCCGTGCCAGCTCGGCCCGCGCAAGCTGGACGCTGGGCAGACCCATGACCTTGCTGACGGTGCGCTCGATGTAATCCGCTTCGTGGCCGGGCTTGCGCTCTGCCAGCGCTGGGCTGGCCTCGGCCATGGCCTTGCCGATTTCCTCGGCACTGCGGCCCCGGCTGGCCAGCTTCTGCGCGGCGATAAAGTCGCACTTGCTGAGGTCATGACCGAAGCGCTTGACCAGCCCGGCCATCTCGCTGCGGTACTCGTCCAGCGCCGTGCGCCGGTGGCGGCTAAGCTGCCGCTCGGGCAGTTCGAGGCTGGCCAGCCTGCGGGCCTTCTCCTGCTGCCGCTGGGCCTGCTCGATCTGCTGGCCAGCCTGCTGCACCAGCGCCGGGCCAGCGGTGGCGGTCTTGCCCTTGGATTCACGCAGCAGCACCCACGGCTGATAACCGGCGCGGGTGGTGTGCTTGTCCTTGCGGTTGGTGAAGCCCGCCAAGCGGCCATAGTGGCGGCTGTCGGCGCTGGCCGGGTCGGCGTCGTACTCGCTGGCCAGCGTCCGGGCAATCTGCCCCCGAAGTTCACCGCCTGCGGCGTCGGCCACCTTGACCCATGCCTGATAGTTCTTCGGGCTGGTTTCCACTACCAGGGCAGGCTCCCGGCCCTCGGCTTTCATGTCATCCAGGTCAAACTCGCTGAGGTCGTCCACCAGCACCAGACCATGCCGCTCCTGCTCGGCGGGCCTGATATACACGTCATTGCCCTGGGCATTCATCCGCTTGAGCCATGGCGTGTTCTGGAGCACTTCGGCGGCTGACCATTCCCGGTTCATCATCTGGCCGGTGGGTGCGTCCCTGACGCCGATATCGAAGCGCTCACAGCCCATGGCCTTGAGCTGTCGGCCTATGGCCTGCAAAGTCCTGTCGTTCTTCATCGGGCCACCAAGCGCAGCCAGATCGAGCCGTCCTCGGTTGTCAGTGGCGTCAGGTCGAGCAAGAGCAACGATGCGATCAGCAGCACCACCGTAGGCATCATGGAAGCCAGCATCACGGTTAGCCATAGCTTCCAGTGCCACCCCCGCGACGCGCTCCGGGCGCTCTGCGCGGCGCTGCTCACCTCGGCGGCTACCTCCCGCAACTCTTTGGCCAGCTCCACCCATGCCGCCCCTGTCTGGCGCTGGGCTTTCAGCCACTCCGCCGCCTGCGCCTCGCTGGCCTGCTTGGTCTGGCTCATGACCTGCCGGGCTTCGTCGGCCAGTGTCGCCATGCTCTGGGCCAGCGGTTCGATCTGCTCCGCTAACTCGTTGATGCCTCTGGATTTCTTCACTCTGTCGATTGCGTTCATGGTCTATTGCCTCCCGGTATTCCTGTAAGTCGATGATCTGGGCGTTGGCGGTGTCGATGTTCAGGGCCACGTCTGCCCGGTCGGTGCGGATGCCCCGGCCTTCCATCTCCACCACGTTCGGCCCCAGGTGAACACCGGGCAGGCGCTCGATGCCCTGCGCCTCAAGTGTTCTGTGGTCAATGCGGGCGTCGTGGCCAGCCCGCTCTAATGCCCGGTTGGCATGGTCGGCCCATGCCTCGCGGGTCTGCTCAAGCCATGCCTTGGGCTTGAGCGCTTCGGTCTTCTGTGCCCCGCCCTTCTCCGGGGTCTTGCCGTTGTACCGCTTGAACCACTGAGCGGCGGGCCGCTCGATGCCGTCATTGATCCGCTCGGAGATCATCAGGTGGCAGTGCGGGTTCTCGCCGCCACCGGCATGGATGGCCAGCGTATACGGCAGGCGCTCGGCACCGGTCAGGTGCTGGGCGAACTCGGACGCCAGCGCCTTCTGCTGGTCGAGGGTCAGCTCGACCGGCAGGGCAAATTCGACCTCCTTGAACAGCCGCCCATTGGCGCGTTCATACAGGTCGGCAGCATCCCAGTAGTCGGCGGGCCGCTCGACGAACTCCGGCATGTGCCCGGATTCGGCGTGCAAGACTTCATCCATGTCGCGGGCATACTTGCCTTCGCGCTGGATGTAGTCGGCCTTGGCCCTGGCCGATTGGCCGCCCGACCTGCTGCCGGTTTTCGCCGTAAGGTGATAAATCGCCATGCTGCCTCGCTGTTGCTTTTGCTTTTCGGCTCCATGCAATGGCCCTCGGAGAGCGCACCGCCCGAAGGGTGGCCGTTAGGCCAGTTTCTCGAAGAGAAACCGGTAAGTGCGCCCTCCCCTACAAAGTAGGGTCGGGATTGCCGCCGCTGTGCCTCCATGATAGCCTACGAGACAGCACATTAACAATGGGGTGTCAAGATGGTTAAGGGGAGCAACAAGGCGGCGGATCGGCTGGCCAAGCTCGAAGAACAACGAGCGCGAATCAATGCCGAAATTCAGCGGGAGCGGGCAAGGGAACAGCAGCAAGAGCGCAAGAACGAAACAAGGCGCAAGGTGCTGGTGGGGGCCATGATTTTGGCCAAGGTGAACAGCAGCGAGTGGCCGGAGGATCGGCTCATGGCGGCAATGGATGCGTACCTTGAACGCGACCACGACCGCGCCTTGTTCGGTCTGCCGCCACGCCAGAAGGATGAGCCGGGCTGAATGATCGACCGAGACAGGCCCTGCGGGGCTGCACACGCGCCCCCACCCTTCGGGTAGGGGGAAAGGCCGCTAAAGCGGCTAAAAGCGCTCCAGCGTATTTCTGCGGGGTTTGGTGTGGGGTTTAGCGGGCTTTGCCCGCCTTTCCCCCTGCCGCGCAGCGGTGGGGCGGTGTGTAGCCTAGCGCAGCGAATAGACCAGCTATCCGGCCTCTGGCCGGGCATATTGGGCAAGGGCAGCAGCGCCCCACAAGGGCGCTGATAACCGCGCCTAGTGGATTATTCTTAGATAATCATGGATGGATTTTTCCAACACCCCGCCAGCCCCCGCCCCTGCTGGGTTTGCAGGTTTGGGGGCGTGACAGTTATTGCAGGGGTTCGTGACAGTTATTGCAGGGGGGCGTGACAGTTATTGCAGGGGTTCGTGACAGTTAGTACGGGAGTGACGGGCACTGGCTGGCAATGTCTAGCAACGGCAGGCATTTCGGCTGAGGGTAAAAGAACTTTCCGCTAAGCGATAGACTGTATGTAAACACAGTATTGCAAGGACGCGGAACATGCCTCATGTGGCGGCCAGGACGGCCAGCCGGGATCGGGATACTGGTCGTTACCAGAGCCACCGACCCGAGCAAACCCTTCTCTATCAGATCGTTGACGAGTATTACCCGGCATTCGCTGCGCTTATGGCAGAGCAGGGAAAGGAATTGCCGGGCTATGTGCAACGGGAATTTGAAGACAATAACT
