## Supplementary material for "The *Coxiella burnetii* QpH1 plasmid is a virulence factor for colonizing bone marrow-derived murine macrophages": S3 Appendix.docx

Complete sequence of the pQGK plasmid:

CTCGAGGGGATAACGCAGGAAAGAACATGTGAGCAAAAGGCCAGCAAAAGGCCAGGAACCGTAAAAAGGCCGCGTTGCTGGCGTTTTTCCATAGGCTCCGCCCCCCTGACGAGCATCACAAAAATCGACGCTCAAGTCAGAGGTGGCGAAACCCGACAGGACTATAAAGATACCAGGCGTTTCCCCCTGGAAGCTCCCTCGTGCGCTCTCCTGTTCCGACCCTGCCGCTTACCGGATACCTGTCCGCCTTTCTCCCTTCGGGAAGCGTGGCGCTTTCTCATAGCTCACGCTGTAGGTATCTCAGTTCGGTGTAGGTCGTTCGCTCCAAGCTGGGCTGTGTGCACGAACCCCCCGTTCAGCCCGACCGCTGCGCCTTATCCGGTAACTATCGTCTTGAGTCCAACCCGGTAAGACACGACTTATCGCCACTGGCAGCAGCCACTGGTAACAGGATTAGCAGAGCGAGGTATGTAGGCGGTGCTACAGAGTTCTTGAAGTGGTGGCCTAACTACGGCTACACTAGAAGAACAGTATTTGGTATCTGCGCTCTGCTGAAGCCAGTTACCTTCGGAAAAAGAGTTGGTAGCTCTTGATCCGGCAAACAAACCACCGCTGGTAGCGGTGGTTTTTTTGTTTGCAAGCAGCAGATTACGCGCAGAAAAAAAGGATCTCAAGAAGATCCTTTGATCTTTTCTACGGGGTCTGACGCTCAGTGGAACGAAAACTCACGTTAAGGGATTTTGGTCATGAGATTATCAAAAAGGATCTTCACCTAGATCCTTTTAAATTAAAAATGAAGTTTTAAATCAATCTAAAGTATATATGAGTAAACTTGGTCTGACAGGGTACCTTATTAGAAGAACTCGTCAAGAAGGCGATAGAAGGCGATGCGCTGCGAATCGGGAGCGGCGATACCGTAAAGCACGAGGAAGCGGTCAGCCCATTCGCCGCCAAGCTCTTCAGCAATATCACGGGTAGCCAACGCTATGTCCTGATAGCGGTCCGCCACACCCAGCCGGCCACAGTCGATGAATCCAGAAAAGCGGCCATTTTCCACCATGATATTCGGCAAGCAGGCATCGCCATGGGTCACGACGAGATCCTCGCCGTCGGGCATGCGCGCCTTGAGCCTGGCGAACAGTTCGGCTGGCGCGAGCCCCTGATGCTCTTCGTCCAGATCATCCTGATCGACAAGACCGGCTTCCATCCGAGTACGTGCTCGCTCGATGCGATGTTTCGCTTGGTGGTCGAATGGGCAGGTAGCCGGATCAAGCGTATGCAGCCGCCGCATTGCATCAGCCATGATGGATACTTTCTCGGCAGGAGCAAGGTGGGATGACAGGAGATCCTGCCCCGGCACTTCGCCCAATAGCAGCCAGTCCCTTCCCGCTTCAGTGACAACGTCGAGCACAGCTGCGCAAGGAACGCCCGTCGTGGCCAGCCACGATAGCCGCGCTGCCTCGTCCTGCAGTTCATTCAGGGCACCGGACAGGTCGGTCTTGACAAAAAGAACCGGGCGCCCCTGCGCTGACAGCCGGAACACGGCGGCATCAGAGCAGCCGATTGTCTGTTGTGCCCAGTCATAGCCGAATAGCCTCTCCACCCAAGCGGCCGGAGAACCTGCGTGCAATCCATCTTGTTCAATCATATGCGCTCTCCTTTCAGAAGGATTAATGTCATTATTTATTTATGGGGTATGGAGAGGGATATTTCAAGGCGCACAAATATGATTCCAGGCTCGTTCTGCGGTTCCGCCTGGGTGTTCGCTGAGATAGGTTTTTACTTCGTTATACGAAAGCGGCGAATCGCCAAAAATTTTTTTTCCAAGTTCGCTGCGAAACGAAGCCATTTCCCGAAGAAGTATGTGCCAAGTGTTTGTAGGTTTTGTTAAAATTTTCTAATAAGGATAATAGTTCATTGCACTTGTCCGTCACAAAGGTTATTTTATCCACAAAAAATTTGCCTACTAGGCCCTCCATTTCAGGAGATAAGAACGCATCATTTATCCATTTGAGTTTTTGGAATGGTAAAGTTTTTAAAAAATCATCTATCTCTGAGGTTTCTTTAGATAGAAGCTTTGGTAGAAAAACATTGTCCGGATTAATTAGAGAACCTGTTTGGATCCTTACTTGTACAGCTCGTCCATGCCGAGAGTGATCCCGGCGGCGGTCACGAACTCCAGCAGGACCATGTGATCGCGCTTCTCGTTGGGGTCTTTGCTCAGGGCGGACTGGGTGCTCAGGTAGTGGTTGTCGGGCAGCAGCACGGGGCCGTCGCCGATGGGGGTGTTCTGCTGGTAGTGGTCGGCGAGCTGCACGCTGCCGTCCTCGATGTTGTGGCGGATCTTGAAGTTCACCTTGATGCCGTTCTTCTGCTTGTCGGCCATGATATAGACGTTGTGGCTGTTGTAGTTGTACTCCAGCTTGTGCCCCAGGATGTTGCCGTCCTCCTTGAAGTCGATGCCCTTCAGCTCGATGCGGTTCACCAGGGTGTCGCCCTCGAACTTCACCTCGGCGCGGGTCTTGTAGTTGCCGTCGTCCTTGAAGAAGATGGTGCGCTCCTGGACGTAGCCTTCGGGCATGGCGGACTTGAAGAAGTCGTGCTGCTTCATGTGGTCGGGGTAGCGGCTGAAGCACTGCACGCCGTAGGTCAGGGTGGTCACGAGGGTGGGCCAGGGCACGGGCAGCTTGCCGGTGGTGCAGATGAACTTCAGGGTCAGCTTGCCGTAGGTGGCATCGCCCTCGCCCTCGCCGGACACGCTGAACTTGTGGCCGTTTACGTCGCCGTCCAGCTCGACCAGGATGGGCACCACCCCGGTGAACAGCTCCTCGCCCTTGCTCACCATGTCAAATCTCCGTTTTCAACTAAAGTTAAAACAAAATCAAAATAAGCCTATATTTAAGCCTATCTTAGCTTTTCTCAAAGCTATAAAATGAAGAATATCGATATCTTTTTTAAATGTCAACACTTAAATCCCATAAAATCATTAAAATAAGAAAAATCTAAGCTTCTACCGATAATTAAGGATATTCAATTACAAAAAAAAAATGCCCATTAATTAAACAATCCGTTTAATTAATGGGCATTTGCTTTCTATTTTGTGTAATTAGGAATCTAATAATAATTCATTTAGCCCTTTCACAAAACTGGCAGGGTCTTTAAGCTGCTCTCCTTCAGCTAGGAGCGCTTGATTAAGCAATAAATCGGCCCAACGATTAAAGCGTGTTTTATCAGATTCATTTTTCACCCGTAAAATTAAGGGGTGCGATGGATTAATTTCTAAAATAGGTTTTGCCTGCATGAAATCCTGACCCGTTTGAATTAATAATCGCTAGCCAACCTCCCTTCTTTCCCAATCCACTGTCCCTGCGAGATCACGTAATCCTTAAATCCTGTAAATCTTTTACTAGTAATTCTAGTTCTTCTACTTCCACTGTTACTCCTCAGCCTTGTCGGTAAAATGCTCGACAAGCCAATCGACTTCTTCCGGCGATGTAAAATGAATAATTACTTTACCTTCCCCTTTCTCATTAATATTGATCGCAATCTTTGAAGACAAGCTTCTGGATAATTGATTGACCCATCCCTGGACCTCGTCAGCATAGGGCGCGGGCTTGGTTTCTTTCGGTGTCTTAGCGAATTGCACTAACTTTTCCGCTTCACGAACTGTTAAATTTTTATCAATAATCTTCTGCGCAAATAAGATTTGTTGATCCTTTGGCAGCGTTAACAATGCTCTCGCGTGGCCCATTTCTAGTTTATCCGTTTGTAACAAAACCTTAACAGAATCGTCAAGAGAAAGCAGTCGCAAAATATTGGTTACAGCGGTACGCGAACGTCCAACTGTTTCCGAAATCGCAGCGTGTGACATAGAGAATTCGTCCCTTAAACGAGAAAAAGCAAGCGCCTCATCAATTGGATTTAGATTTTCCCGTTGTATATTTTCAATAAGCGCAAACGCTAATGCTGTAGTATCATCAACATTTCTGATAATGGCAGGTACTTTTTTTAACCCCGCTTCTTTCGCTGCGCGCCAGCGTCTTTCTCCCGCAATGATTTCATATCGATTGGTTTGAATTTGACGTACAATGAGTGGTTGTATAATGCCTTGAGATTTAATTGAATTCACAAGCTCCTGTAAAGATTCTTTTATCCAACCCTTTCGGGGCTGATATTTACCCGTCTGTAGTAAATCAATGGAGAGAAAATATAAATTTTGAGCAACGAAATCGTTTGTTTTAGAGGCGTTCGCTAATGACTTCAAATCCCTTAGACCTTTTAAGCCTTTCAATTCTATATCAGCCATTATTTTTCCTTTCAATACATTTACGTTATCGCGTACCGCAAACGTCTCCTCCATCCTGCAAATGTCTCAACCAGCAAGAACTCCTTTCATTTTCAATACCGCAAACGTCTCGCTTTATCCTGCAAATGTCGCATATCAACCCGCAAACGTCGCTTCCTATCCTGCTAATGTCTCTTAAAAGTTTTCAAGATCATGAAAATTAAGCAATTAATAAATTGTAAACAACAGAAACTAATTAAAAAAATTAAAACACGTTGTGTTTCTTTTGTAAATAGAAACTTCATAAAAAAGTTGCATTTTCATAAAAAACCAGATATTTTTTAAAAAGGATGAAATAATTCATAGTTTACAAAGGAACTCGTCATGCTAGAAACACAAATTACCCCCTACGGTACAGAAACCCCCGAACAATTAATGGACAAATTCTACCAAGCTGGTAACGAAATGCTATTAACACTTCGAAATTATATTACTAGCCCAGATAAAAGAAAAAAATCTCGGACGTGGGGTGCGATAGAAGCTGCGAAAATGGTTGGAGTCTCCGCGCCTACTTTCAGAAAACTATTGGAATCCGATAACGAAGTTCCTGGAATCATAATTGAAGAAAATGAGAACGGAAGGAAAATAAAAAAGTACACGCTTACTGCGATTAACAATTTGCGTGAAAAGGCAAAAACCCGTTACAAGCGCCCTAAAGGCTCTAAACCATTAACAATAGCAATATCGAATTTAAAGGGCGGCGTCGGGAAAACAGAAACGGCAGTTGATTTAGGTAAAAAAATAGCCATTGAAGGCTTACGTTCCCTCCTTCTCGACTTTGATGCTCAAGGCACAGCAACCCTTATAAGCTCAGGACTGATCCCTGACTTAGAATTACGTTACGAAGACACGATTACCAACACTTTAATTTCAGATCCGAACAACATTAAGAATATCGTTCTTAAAACACATTTTGATGGTTTCGATATTATTCCTGCCAATCTAGCGATACAAGATTGCGATTTAATCCTCCCAAACGATAAAGAAAACAACAACGACCGTTTAGGTTCTCCTTTTCTAAGGTTGGCAGAGTCTCTAAAAATAATTAAAAATCAATATGATGTAATACTTATTGATTGTGGTCCTAATTTGGGATTACTCACATTAAACGCCATTATTGCTTGTGATGGCATGATTATTCCTATTCCGCCCAGCATGAATGATTACTCAAGCTTCATCATGTACACAGCAACTTTACGTAATATGTTTAGAGAATTATCGAACAAAAAGTTGGACTATTTGAGAATACTTCTATCAAAGCACAACAGCAGCAATGAGGCTCTGCAAATGGAAAATATGATGCGCGAGCAATTTGGCCGGTATATTCTTTCTAACCATATGTGTGAGACTGTTGAAGTATCGAAGGCGGCAAATGAAATAGGAACGATCTATGATGTATCAAAACCACGAGGAAGTCGAGAGGCTTATCGTAGAGCCCTTCAACACCTTGACGATGTCAACATGGAAATAATTAATAATTTTAAAGACATATGGAAAAGCCAAGTAAAAGTGCTAACAACTTTAGGAGAAACCGTCAATGGATAACAGCAAGCGAAACATTCATAACTCTGGACCGTTAGGTATGTTAATGAAAAATGGCCAGATTAAAAAAATTGAAAATTCAGCAGAATCTAACGAAGGTACCGTCGTATTAAATAAAGCAGCACCCTCCTATTTCAAAACACAAGCGGGGATTGAGTTTACCGAACATGAATTGATCTTTGTGGATCCAAAAGAATGTGAGCCCTGGGAGTATGCTAATAGACAAGATGAGGAACTTGGCAATATAAACGAATTAATCGAATCAATTAAATCAAATAAACAACTACAGCCAGCCCTTATTCGGAAACACCCTCACCCACATGATGACGTTAAATATGAAATTATTTTCGGTCGTCGACGACATATAGCCTGTCTTAACCTTGGTATCCCATTTTTAGCGATTCTCAAAGAGATTCCTAATGTTCAAGATGCAATCGCTTTTCAAGACGCAGAAAATAAACTTAGGAACGACGTCAGCAATTATTCAAACGCCATACTATACAAACGTTTGATAGAAGAAGGGGTTTTCAAAAAGGAAAAAGACCTCGCCGAGAAATTGCGATTATCGCCTTCTACCTTGAATGATTTGATGGCGTATACAAAGATACCAAGTGCTATTGTCAAAAAAATTCCAAACATCCATGCGTTGTCCAAAAGCATTGTGCTTAAAATAGTTCAATTACTCAATAAATCTTCAAAAAATCACGCAAAACTTATTGCTATTGCACCAGATATTGGTAAGTCCATTACTTCACCGGCTAAACTTGAAAGTGCTGTCGAAAAACCAGTAGGCAGCAAAACAAAACAGCGGTTGCAGGCAACTAAACAATATAAGACTAAAGATGGGAAAAAGCTTTTTACGTTTAAAATTGATCATCGAGGCGCGCCTTGCATTGTTTTGAATAAAGAAATCCTCAACAGGGTAGATATGGATACCATGTGTGAGAAAATTAAAAGCCAACTAGAAATTGAATTAAGTCAATCCGGAGCTCCGGATTGACGCCCAATCCGGAGCTCCCGGATTGGAGCTTGAGCCAAATATTGATGGAGAAACGAATGAATTTGGCTGTGGCTACGGACAAAAGAGTCGAACTGAAAAAGCATGTCAATGCTATTCATTGTTCAAATAACCTTACTTTGGTGCAAAGAAAGTTATTCAATGCACTTTTATTTAATGCTTATCCAGAGTTACCCTACAAGCTCAAATTTCAAATTTCAGCAAAAGATTTATGCAAATTAATAGGATATAACAGCAATGACTATGGAAAGCTTAAAAGGGCGCTACTTGGGTTAATCACAACCGCAATTGAATGGAATGTAATCGATTGTGACACAGGGAAGGAAAAAAATTGGAAAGCAAGTTCAATCCTATCAGCGGCTCAGCTGGCGGAAGGCGTTTGTACTTATGAGTACAGTCATATTATGAAAGAATTGCTTTTCCAGCCGGAAATTTATGGAAGAATAGATGTAAAAACCATGTCAAAATTTAAATCAAGCTATGGTTTAGCATTATATGAAAACTGTATTCGCTACCAGAGACTGTCACAGACACCTTGGTTTCCTTTAGATGTTTTTAGAAAACTCATGGGAGTTTTAGGCGGTAAATACACTTCATTTAAAGATTTTAAAAAAAGAGTACTCAATATTGCAGTGAATGAGGTAAATAACCTATCACAAATTCAGGTCTTACCAGAAATCGAGAGACAAAACCAAAAAGTCACAAAGATTAGATTTAAACTGAATAAAAAACACCTTGCGTCATCCGAAAAAATCTCAAACCTAATTAATGCAGAGTTAGAAGAAACCTTAACAAATACATTTAGTCTATCTCCGCTTCTTATTCGAGATTTGCTTACAGAGTATGATATAGCGTTCATTCGTGAAAAAGTGAACATCATTATAAATTCCAGTAGTTTTCAAGAAGGGAAGATACGCGGCCTTAGTGGGTATTTAATAGACTCATTACGAAAAGATTATAAAGCCAGCAAGTCGAGCCAGATGCTTATTAGTGAAGCACGAATGAAGAGGAAATTAGAAGAAGAAGCTGAGAAAGAAAGGGAAGAAAGACGAAGAAATAGATATGAAAAATACGTTAAGGATAAAATAAACAAATATCTTACATTACTAACGGATGCTGAGAAGGATGCTCTAGTGATGGATTTTGAGGTTGAATTGAAGTCAAATAAAAGTAATAAAGTCTTTTACTCTTGGTATAAGAAAAATGGTTTTGAGCATGTTGGGGTTAAAGCTTGCTTTCACAATTTTGTGAAAGAGCATAAAAAACAGCATATGGGAGGAATTCTGTCATTTGAAGAGTTTATAAGCTTGGTGGATGAATACACTTAATAGACAGGCGCGACGTTTGCGGGACGGGAAATTCAGATTCGCGAGGGGGGACGTTTGCAGGATGTTCTCTTTGGAGACTTTATCGAAATATTAATCCTCAAAAGTTACATCTTGTTCTTTCTTCAAACGACTTTTAAGTTCTTCATTAGAATTACTAGCGGCTTCATAAAGCTTGAGGTATTGTTCCGCGCGTCGTTTATATTCTTTAACCGCGAGCTGAGCTTGCAAACCGGTCTGGTAAATTTTAGCCAGTATCGCTTTCAATTGCTGTGGATTATCAGGCAGCGAACGAATCGCCTCGGGAAGGGGTGTTGGTGCGATATTTTCGGATTGAAAAAACTGAGGGGAAAGCGTTTTAACTTCCTCTTTGAGCCGTTCAATTTCCTCTTTGAGAGTAAAAAATTCGTGCAAAGTACCTTGGGTGATTTCCTCAAAGATTCGTGCACGCGCTTTGGCCATGCGTTGACGCTCATTCCAGGGCTCTGGGTATTCTTCAGCCGCGGCTTGAACCATCGCTTCAAATCGTTTTTTGGCGTCCGCGCGCACTTGGAGCTCGAAACGAACCAATCCTTTGTTGGCTTGCGAACGCCGGTATTTTTTTTGGGCATTGGGTTTGTGGCGCGTCGGAATTGATTTGATTTTATCCATACCATTTTTATCCTTTTTCACCGGAGATAATATAGCAGTTTTTAGATCGGTTGAGGAGGACAGCGATTCTACAAAGAAT
