## Supplementary material for "The *Coxiella burnetii* QpH1 plasmid is a virulence factor for colonizing bone marrow-derived murine macrophages": S4 Table.docx

**S2 Table. Primers for PCR Identification**

| **Primers** | **Sequences（5’-3’）** | **Note (area)** |
| --- | --- | --- |
| cbua-01F  cbua-01R | TGAAAGACAGTCTCATTACCT  GAAATGTTTCCAGGACTTGC | CBUA0001 |
| cbua-03F  cbua-03R | TTGGTGAGTCGTCAGTATTTTG  TTACACATTTTTAATACAAGC | CBUA0003 |
| cbua-05F  cbua-05R | TAAGAACACGTTAGTAGAAGAG  GTTTTAGAACGTGGCAAGGT | CBUA0005 |
| cbua-06F  cbua-06R | TGTTTGCAACTGTTCCATCCT  TGATGTTGATGGCGCAGACTC | CBUA0006 |
| cbua-07F  cbua-07R | TTGCAGGGACTGTGTGTAACG  CTAACTTCTATTGGCTCTCTTC | CBUA0007 |
| cbua-08F  cbua-08R | ATGAATAATAACTTGGAAAATG  TTCGCCACTTTTCCCTGTAAT | CBUA0008 |
| cbua-08aF  cbua-08aR | AATCAAGTTGCTATTAGGGTGT  CATCTTCATTCCACGCCTTAC | CBUA0008a |
| cbua-08bF  cbua-08bR | TCAATAAGTCGTGAGCAATCC  ATCGTAGCGTTGCGTCACAC | CBUA0008b |
| cbua-08cF  cbua-08cR | GGAATTAACCCTTATGCGTAT  CGTTAGCGCCTCGTCATTAC | CBUA0008c |
| cbua-10F  cbua-10R | TTGATGCCGTTCTCTTAGTTC  CTATTGAAAATATCACTGCTGG | CBUA00010 |
| cbua-11F  cbua-12R | AGTGATATTGGCACAACGATG  AACGTTCCTCGGCAAGACT | CBUA0011～CBUA0012 |
| cbua-13F  cbua-13R | ATGCCATATTTTTTTACACTAC  GAAAACAGTTGTTATTAGTG | CBUA0013 |
| cbua-13aF  cbua-14R | TAATTTATTAGGGTGCTTGC  GCTCAAATTCCGCACCTACT | CBUA0013a～CBUA0014 |
| cbua-15F  cbua-15R | TTTGAGCGATGTAAATGACTAT  ATTCCTCTTCAGTTTCCGTTTC | CBUA0015 |
| cbua-16F  cbua-16R | ATGAGATTAGAACAACCAAG  TGTTCATTTTCTGAGTCCGAG | CBUA0016 |
| cbua-17F  cbua-17R | GAGAAAAAATGTCACGAGAG  CCCTTTCCTCTATGCGTTCT | CBUA0017 |
| cbua-18F  cbua-18R | AGGATACGATGGTTCTTGTG  GAAGCATGAAAGCAAAGTATC | CBUA0018 |
| cbua-20F  cbua-20R | ATCGATTTTTCGTGGTTCTC  TCTTGAAGCTTACCAAAATGC | CBUA0020 |
| cbua-21F  cbua-21R | GTTTCTTTGCATTGCTGTTG  GAAGTATATGGGTATCAAAGTG | CBUA0021 |
| cbua-22F  cbua-22R | CTACAATGCGCCTATGGAC  CTATGCGTTATCCACTGTTTC | CBUA0022 |
| cbua-23F  cbua-23R | CCCTCATTTTGGTATTTCG  CAATTGTGCTATTGAGGAAAC | CBUA0023 |
| cbua-24F  cbua-24R | AGCAGCGCATTTAATCTTCT  GTGGGGCACATCTACTTTC | CBUA0024 |
| cbua-25F  cbua-25R | GAAATTCTTTCCTCTTTGTAGC  TTGAATAAGCCAAATGATG | CBUA0025 |
| cbua-26F  cbua-26R | TGATTGTCTATCGAAGCCTAG  GCATGGGAGAATGAGCAGTC | CBUA0026 |
| cbua-27F  cbua-27R | GAAGGCCTTTTCTATCAACG  TTAAACCTTTATCGCCCATTG | CBUA0027 |
| cbua-28F  cbua-28R | CACCTCTATCAGTTCGCCTAT  TGACTGATGGAAATCTGCTC | CBUA0028 |
| cbua-29F  cbua-29R | CACCCAAACAGCTCACGATC  CGTGGGAAATCGGTGAGGT | CBUA0029 |
| cbua-29aF  cbua-29aR | CAGCAAAATTAAGATGGACT  GGAATATCCGTGATTGCCTC | CBUA0029a |
| cbua-31F  cbua-31R | GTCAATTTTTGGATTTCACATAG  TTATCTTCCATACAAATTCATC | CBUA0031 |
| cbua-32F  cbua-32R | ATGGAAGATAACTGCATTAAAAT  TTATTTTATCCAAATACAGTTAG | CBUA0032 |
| cbua-33F  cbua-33R | ATGAATCCTAAAGATTTACTCTAT  AATTATCTTTCCCCCCTTCTAC | CBUA0033 |
| cbua-34F  cbua-34R | TCAACATTTTATCATCAGAGG  TTGGGCGCTTACTTTAGGC | CBUA0034 |
| cbua-34aF  cbua-34aR | TTCACTCGCTGAAAAACCTTC  AAAGTGGGCAAATTGGTTCG | CBUA0034a |
| cbua-36F  cbua-36R | TGTTACTCCTCAGCCTTGTCG  GTACGCGATAACGTAAATGT | CBUA0036 |
| cbua-37F  cbua-37R | AAACACAAATTACCCCCTACG  GTTATCCATTGACGGTTTCTCC | CBUA0037 |
| cbua-38F  cbua-38R | AGCGAAACATTCATAACTCTG  TGACTTAATTCAATTTCTAGTTG | CBUA0038 |
| cbua-39F  cbua-39R | CCAAATATTGATGGAGAAACG  AGTGTATTCATCCACCAAGC | CBUA0039 |
| cbua-39aF  cbua-39aR | TTAATCCTCAAAAGTTACATCT  TAAAATCAAATCAATTCCGAC | CBUA0039a |
